# Linking ecological specialization to its macroevolutionary consequences: An example with passerine nest type

**DOI:** 10.1101/2021.08.24.457563

**Authors:** Rosana Zenil-Ferguson, Jay P. McEntee, J. Gordon Burleigh, Renée A. Duckworth

## Abstract

A long-standing hypothesis in evolutionary biology is that the evolution of resource specialization can lead to an evolutionary dead end, where specialists have low diversification rates and limited ability to evolve into generalists. In recent years, advances in comparative methods investigating trait-based differences associated with diversification have enabled more robust tests of this idea and have found mixed support. We test the evolutionary dead end hypothesis by estimating net diversification rate differences associated with nest type specialization among 3,224 species of passerine birds. In particular, we test whether the adoption of hole-nesting, a nest type specialization that decreases predation, results in reduced diversification rates relative to nesting outside of holes. Further, we examine whether evolutionary transitions to the specialist hole-nesting state have been more frequent than transitions out of hole-nesting. Using diversification models that accounted for background rate heterogeneity and different extinction rate scenarios, we found that hole-nesting specialization was not associated with diversification rate differences. Furthermore, contrary to the assumption that specialists rarely evolve into generalists, we found that transitions out of hole-nesting occur more frequently than transitions into hole-nesting. These results suggest that interspecific competition may limit adoption of hole-nesting, but that such competition does not result in limited diversification of hole-nesters. In conjunction with other recent studies using robust comparative methods, our results add to growing evidence that evolutionary dead ends are not a typical outcome of resource specialization.

---

Resource specialization, where species use a narrower range of resources compared to related taxa (Futuyma and Moreno 1988), is a common phenomenon in evolution. There are many reasons resource specialization (hereafter specialization) can evolve. For example, specialists can benefit from reduced competition, or from the avoidance of predators or parasites (Futuyma and Moreno 1988; Bernays 1989; Schluter 2000). Once evolved, specialization can have diverse consequences for macroevolutionary dynamics. Historically, specialization often has been considered an evolutionary dead end, possibly resulting in both reduced net diversification relative to generalists, and difficulty evolving from specialist to generalist (Day et al. 2006). This difficulty is thought to arise when adaptation to specialized resources occurs across numerous traits (Futuyma and Moreno 1988), and, when coupled with changes in resource availability, leads to elevated extinction rates in specialists. Additionally, reduced diversification of specialists could result if the number of available niches in the specialized state is low.

Consistent with the evolutionary dead end view of specialists, the prevailing historical viewpoint was that specialists evolve from generalist ancestors (Schluter 2000). Prior studies have found mixed support for this viewpoint, with inferences from phylogenetic comparative methods showing that generalists regularly evolve from specialists in many clades (Day et al. 2016; Sexton et al. 2017; Villastrigo et al. 2021). These results indicate that the dominant direction of evolutionary transitions involving specialization should be treated as an open question for any specialization scenario.

More recent studies have also noted the potential for specialization to increase diversification through multiple mechanisms. First, the release of specialists from the effects of competition and/or predation could trigger periods of niche-filling diversification (Schluter 2000), leading to higher rates of speciation in specialists compared to generalists. Further, in circumstances where specialization releases specialists from competition or predation (Futuyma and Moreno 1988), specialists may experience greater population persistence than generalists, which could increase diversification rates by decreasing extinction rates or by increasing rates of allopatric speciation via the longer survival of incipient lineages (Harvey et al. 2019), or both. Finally, specialist lineages could also have higher diversification rates if more specialized lineages have more fragmented distributions or lower rates of dispersal (Gavrilets et al. 2000; Birand et al. 2012).

Recent diversification models and statistical developments have the potential to alter our conclusions on the macroevolutionary consequences of specialization. A growing awareness of type I error (Davis et al. 2013; Rabosky and Goldberg 2015), the importance of large sample sizes for statistical power (Davis et al. 2013; Day et al.2016), and the misspecification of the null hypothesis for the original state-dependent diversification models (Beaulieu and O’Meara 2016; Rabosky and Goldberg 2017; Caetano et al. 2018) have produced a wave of new studies using comprehensive statistical approaches on the question of whether specialization is linked to diversification. The majority of studies after Day et al.’s (2016) review on specialization have found either no association between specialization and diversification (e.g., Alhajeri and Steppan 2018; Crouch and Ricklefs 2019; Villastrigo et al. 2020) or have found that specialization is associated with higher diversification rates (e.g., Conway and Olsen 2019; Otero et al. 2019; Tonini et al. 2020), with few studies indicating specialization leads to an evolutionary dead end (e.g., Cyriac and Kodandaramaiah 2018; Day et al. 2016). While these newer studies have relatively large sample sizes (compared to pre-2016 studies) and more transitions to specialist states, some of the sample sizes are still too small to fit state-dependent diversification models with more than two states (Davis et al. 2013) and to have enough power to estimate specialization’s consequences for diversification (Davis et al. 2013). This has also been difficult to accomplish because, for the majority of the software that fits state-dependent diversification models, estimating intervals requires complex likelihood function maximizations (Zenil-Ferguson et al. 2018) or many bootstrap simulations to approximate confidence-likelihood intervals for diversification parameters (FitzJohn 2012; Beaulieu et al. 2021). Moving forward it is necessary to integrate models that incorporate more states and more parameters that mathematically link multiple specialist and generalist states to the process of diversification while accounting for the potential for heterogeneity in the diversification process (Beaulieu and O’Meara 2016). Furthermore, the calculation of uncertainty for the parameter estimates in more complex models is key to understanding the power of the sample and testing whether specialization is linked to diversification.

Here, we highlight these methodological approaches with a large dataset on nest type for 3,224 species of passerine birds. Passerines provide a unique opportunity to examine the macroevolutionary consequences of specialization. Specifically, hole-nesting is a prime example of resource specialization which has evolved multiple times across the clade of 6000 passerine taxa, providing the power needed to estimate rates of diversification linked to trait evolution. Second, passerines can be grouped into three general nesting habits: open-cup nesters, dome nesters and hole nesters (e.g., Wallace 1868; Martin 1995; Collias 1997). Open-cup and dome nesters are relatively unspecialized in their nesting type compared to hole-nesters; however, similar to hole nesters, dome nesters have reduced predation rates compared to open-cup nesters (Oniki 1979a; Linder and Bollinger 1995; Auer et al. 2007; Martin et al. 2017).

Therefore, our approach of comparing adoption of these three nest types enables us to simultaneously assess the potential effects of nest type specialization and escape from predation on diversification rates (Fig. 1).

**Figure 1.**
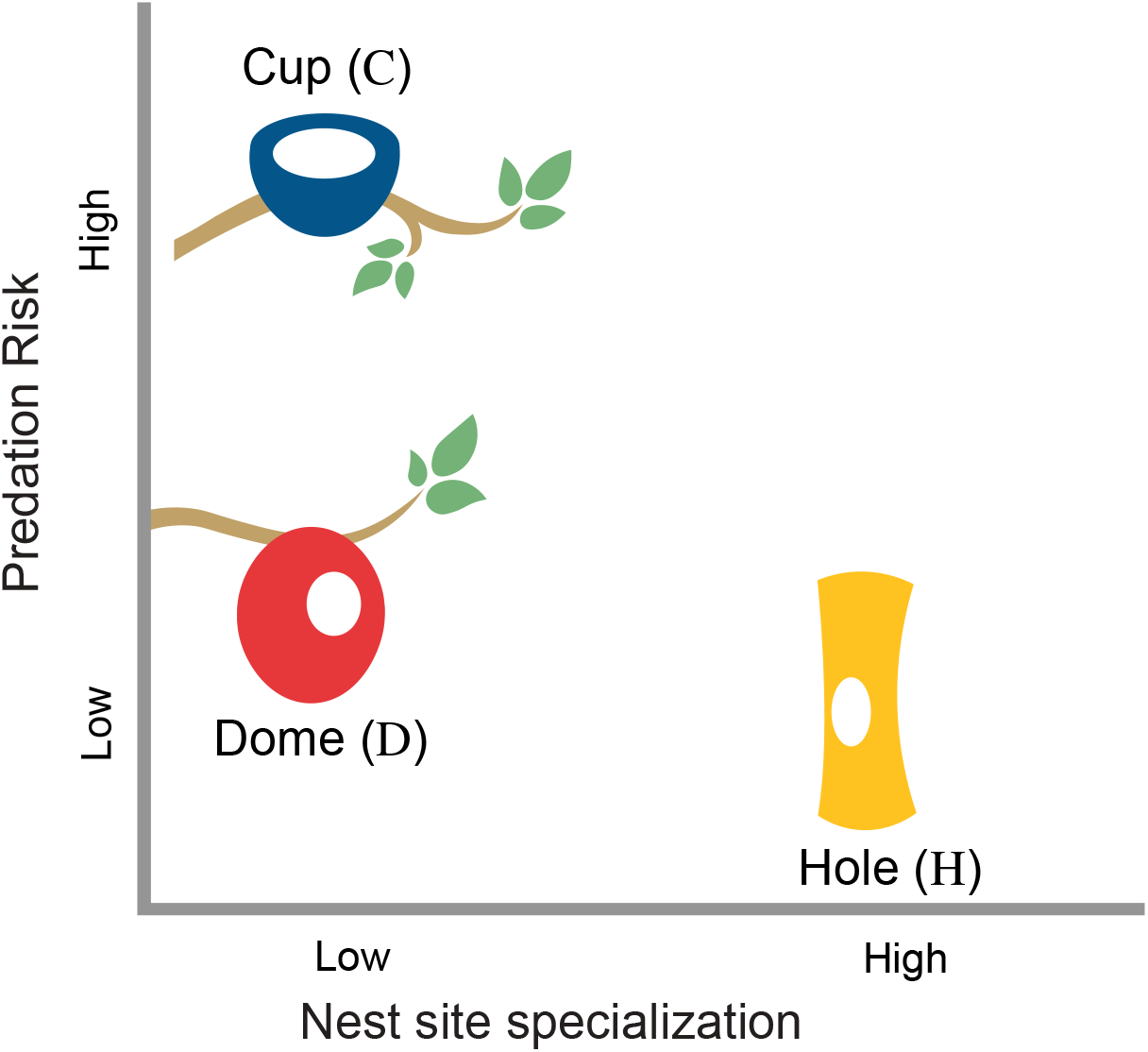
The nest types of passerine taxa differ by how specialized they are in nest site and in their predation risk. Hole is the most specialized nest type and has the smallest predation risk, followed by dome with also a lower risk but relatively unspecialized in its nesting substrate. Finally, open-cups have the highest predation risk but are also considered unspecialized. Nest colors represent the colors of the states for the models presented.

We employ a graphical modeling approach (Jordan 2004) in Bayesian statistics to infer point estimates and credible intervals of parameters from state-dependent speciation and extinction models (SSE models), and diversification-free models (Mkn-Markov models without state-specific diversification parameters) coded in RevBayes software (Höhna et al. 2016). The large nest type dataset and phylogenetic tree are the input to models that are estimated in a Bayesian framework and this enables us to avoid a number of the common pitfalls discussed above. Additionally, we adopt a systematic approach to specifying prior distributions to model enhanced extinction rates to assess the consistency of the statistical inference across a spectrum of macroevolutionary diversification scenarios. Finally, after determining the best model for the evolution of nest type, we used it to reconstruct ancestral states of nest types to visualize the pattern of evolutionary history for the trait and compare with previous ancestral state reconstructions.

## Nest type evolution and its potential links to diversification

Nest type selection is a critically important aspect of avian habitat because nest failure rates are high in birds (Nice 1957) with predation as the main cause of nestling mortality (Ricklefs 1969; Martin 1993). Adoption of hole-nesting has the advantage of protecting offspring from predation, as non-excavating hole nesters show an approximate 43% reduction in nest failure compared to non-hole nesters (Martin 1995). Thus, a major ecological consequence of hole-nesting specialization is that it increases nesting success by providing a release from predation. Hole-nesting represents resource specialization for two reasons: the great majority of species that nest in holes do so obligately, and exclusive use of holes greatly restricts the substrates that are suitable for nest building.

Hole-nesting is associated with the evolution of a suite of traits, such as increased nesting period, brighter egg coloration, and larger clutch sizes, that may make it difficult to transition out of hole-nesting (Martin and Li 1992; Kilner 2006). Moreover, the great majority of hole-nesting passerines cannot excavate their own nest holes, and competition for this limited resource is intense (Newton 1994). While hole-nesting likely evolves as a response to predation pressure, the intense competition for nesting sites decreases nest hole availability (Cockle et al. 2011). This competition may limit ecological opportunity for hole-nesting lineages despite the reduction in predation pressure. Further, reliance on holes for nesting may increase extinction risk when environmental changes reduce nest hole availability, particularly if transitioning from hole-nesting is difficult. This combination of characteristics makes hole-nesting in passerines a form of specialization that could be an evolutionary dead end. However, as hole-nesting decreases nest predation, it is also possible that adopting hole-nesting could lead to greater population persistence over time which, in turn, could lead to reduced extinction rates or increased rates of allopatric speciation (Harvey et al. 2019). Thus, it is also possible that hole-nesting lineages could experience greater diversification rates than the more generalist non hole-nesting lineages.

The evolutionary dead end hypothesis assumes that it is easy to adopt hole-nesting habits, but difficult to transition out of them because of secondary adaptations that restrict evolutionary transitions (Collias and Collias 1984). In passerines, most hole nesters are secondary (i.e., they cannot make new cavities, and thus they rely on what is already in the environment), and so adoption of hole-nesting is mainly a behavioral shift not requiring extensive morphological modification (although see above for examples of life history traits associated with hole-nesting). This could make it relatively easy to evolve hole-nesting behavior. However, the intensely competitive environment that hole nesters face could mean that hole-nesting niches are already saturated, making it a more difficult habit to adopt. To examine these possibilities, we not only assess the impact of nest type on diversification rates, but we also calculate transition rates between states.

## Materials and Methods

### Data

Our nesting data is a slightly adapted version of a dataset originally assembled for McEntee et al. (2021). This dataset was generated by scoring nesting behavior of passerine species using descriptions from the Handbook of the Birds of the World Alive, (Del Hoyo et al. 2017, last accessed 30 June 2016, hereafter HBW). Specifically, species’ nests were scored as either open cup, dome, or hole. A few species show flexibility in nest type use among these three types, and are capable of nesting in, for example, either an open cup or a hole. We scored these examples accordingly as nesting in either. “Hole” refers to any nest built inside a tree cavity, rock crevice, or earthen bank. The small number of passerine brood parasites were excluded from analyses. (McEntee et al. 2021) were able to score the nest types for approximately three quarters of the 5,912 passerine species from the information in the HBW species descriptions.

Next, we matched the passerine species in the nesting dataset to the species’ names on the tips of the avian supermatrix phylogeny of Burleigh et al. (2015), one of the largest phylogenetic trees of birds constructed exclusively from molecular data. This tree was time-calibrated using penalized likelihood in r8s (Sanderson 2003) with twenty fossil calibrations from throughout the avian phylogeny (Baiser et al. 2018). We identified all cases in which a species in the Burleigh et al. (2015) phylogenetic tree, which used the Clements taxonomy, did not have a corresponding species with the exact same name in the dataset from the HBW Alive/BirdLife International taxonomy.

This was due to either missing nesting data or a taxonomic mismatch. For taxonomic mismatches, we examined the taxonomic history for these species in Avibase (https://avibase.bsc-eoc.org/), and when appropriate, changed the species name in the nest dataset to match the phylogenetic tree (Burleigh et al. 2015). Taxa treated as subspecies in the HBW taxonomy (2015) but as species in Burleigh et al. (2015) were not included in our analyses because they did not occur as tips in the phylogenetic tree. We then trimmed the phylogenetic tree to include only species in the nesting dataset from HBW. The resulting data set has the nesting state for 3,224 passerine species. Of these, 1943 species had cup nests (*D*), 722 had dome nests (*C*), and 458 had hole nests (*H*). Among species with multiple nest types, 60 were cup or hole nesters (*CH*), 29 were dome or cup nesters (*DC*), and 13 were dome or hole nesters (*DH*). In the few instances (less than 1% of internal nodes) where this tree was not bifurcating (required for the diversification analyses described below) because nodes were collapsed in the r8s analysis, we resolved bifurcations randomly using the function `multi2di` from the R package `ape` (Paradis and Schliep 2019).

### Modeling nest type linked to diversification

Using the three states assigned to the tips of the 3,225-taxon phylogenetic tree (3,224 passerines plus a single species representing the parrot outgroup), we defined four state-dependent speciation and extinction models (SSEs) and one Markov model that we call diversification-free (because it does not contain speciation and extinction parameters Fig. 2). The first multistate SSE model, called MuSSE-3 (Multistate speciation and extinction with three states, Fig. 2a), uses three main states: dome *D*, cup *C*, and hole *H* (Fig. 2a), each with their own speciation and extinction rate. MuSSE-3 also has transition rates between each state (*q*_*ij*_ with *i, j* = *D, C, H*, and *i* ≠ *j*) governing the rate of evolution from one nest type to another. All transitions between nest types are possible. Taxa that have multiple nest types are coded in the input data as belonging to multiple states simultaneously. For example, a bird taxon reported to have nested in both hole and dome nests is coded in the input data as (*H, D*). For taxa (tips) with two nest types, the likelihood calculation includes the product of two transition probabilities from the most recent common ancestor to each potential state in the tips instead of a single transition probability from the most recent common ancestor to one single state.

**Figure 2.**
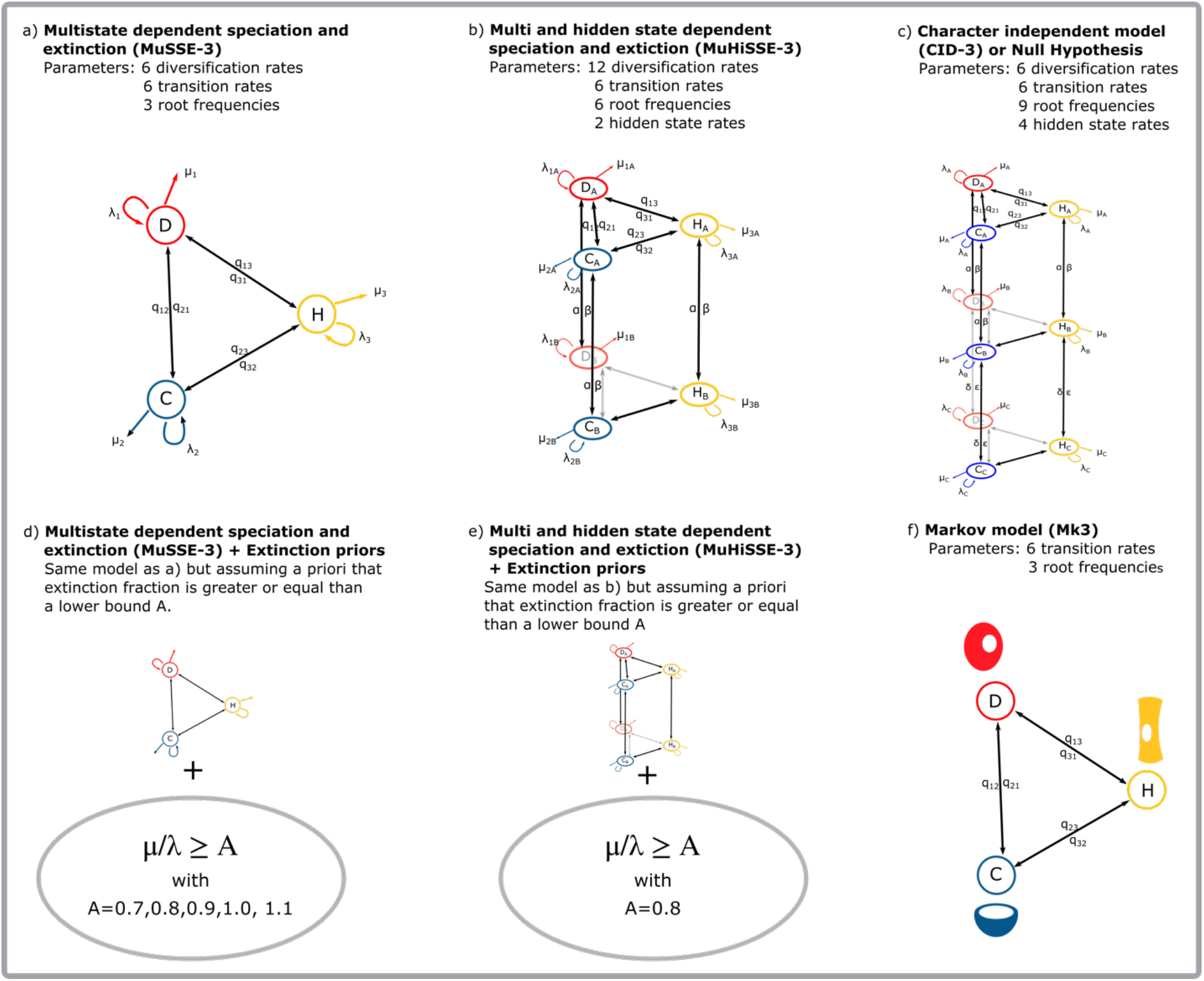
Models of nest type evolution. a) The multistate dependent speciation and extinction (MuSSE-3) assumes transitions among each nest type (*D, C, H*) and diversification rates (speciation and extinction) associated with each state. b) The multi- and hidden state dependent speciation and extinction model (MuHiSSE-3) expands the model to consider hidden states A and B (and transitions between them) to model the potential of other factors along with nest type in the process of diversification. c) The character independent model (CID-3) allows us to assess whether diversification is linked to some unmeasured factors that are independent from nest type. d) MuSSE-3 with prior distributions that restrict the parametric space allows us to infer trait-associated diversification differences where relative extinction is enhanced, a part of parameter space that is typically poorly explored in SSE models. e) MuHiSSE-3 with prior distributions that restrict the parametric space allows us to infer trait-associated diversification differences where relative extinction is enhanced. f) The Markov model (Mk3) assumes that there are no diversification rate differences among nest types, and transitions among nest types (*D, C, H*) are allowed.

The second model (Fig. 2b) is the hidden state extension of the MuSSE-3 and it is called multi and hidden state dependent speciation and extinction model (MuHiSSE-3). The hidden state extension includes background heterogeneity in the diversification rate that is not linked to the trait of interest (Beaulieu and O’Meara 2016), and (Caetano et al. 2018). The MuHiSSE-3 model expanded from three single nest type states to six with subscripts *A* or *B* (Fig. 2b).

The character independent model (CID-3) or null hypothesis model (Fig. 2c) accommodates heterogeneity in the process of diversification completely unlinked from nest type but due to some unmeasured factors called hidden states. CID-3 specifies a null hypothesis that is comparable to MuSSE-3 (Beaulieu and O’Meara 2016; Caetano et al. 2018) because both models have the same number of diversification rates. That is, the CID-3 has six speciation and extinction parameters, but makes the net diversification for main nest type states *D, C, H* equal while assuming that the source of differences in diversification come from the hidden states *A, B, C*. In mathematical terms the null hypothesis (*H*_0_ or CID-3 model) for a state-dependent diversification model with three states and six diversification rate parameters is:

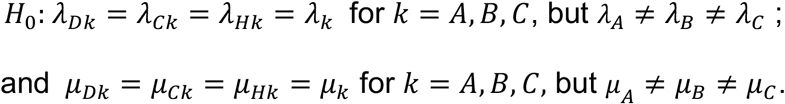

Note that in the CID-3 notation the 3 refers to the number of hidden states that have different diversification rates, and not necessarily the number of states of the main trait as discussed in Caetano et al. (2018).

For all SSE models in Fig. 2 we defined a prior distribution for the log-transformed speciation rates *λ*_*i*_ to be a normal distribution with hyperparameters 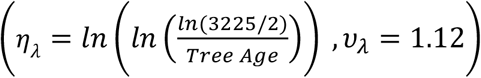, with the phylogenetic tree age as 64.51 million years old. This choice of hyperparameter *η*_*λ*_ for the log-transformed speciation rate makes the expected average rate for speciation approximately 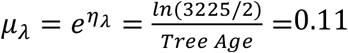, representing an approximation to the current observed extant diversity of 3225 taxa in our tree conditional on having two extant lineages at the root of a phylogeny that is 64.5 million years old (FitzJohn et al. 2009; Freyman and Höhna 2019; Zenil-Ferguson et al. 2019). Note that the age of the passerine crown in this phylogeny is 40.53 my but when adding the *Strigops* (parrot) outgroup as we have the resulting phylogeny is older. Defining hyperparameters and prior distributions is subjective, a broader discussion on eliciting priors and their hyperparameters was added to supplementary material Fig. S14 and S15, showing that inferences are consistent and our likelihood function is informative for SSE models with the data presented. The same prior distributions were used for log-transformed extinction rates. For all models, we assume a sampling fraction *ρ* = 0.51 since our tree contains about 51% of all passerine taxa.

In the supplementary information we fit state-dependent diversification models with six states including taxa with two types of nesting as a separate state (i.e., cup and dome nesters get their own state *DC*) to investigate if different state-coding changes diversification and/or transition rates (Fig. S5).

### Testing the role of diversification in nest type evolution

To test if nest type evolution is linked to the diversification of passerines, we calculated the marginal log-likelihoods for the MuSSE-3 and the character independent model (CID-3). Using the log-likelihoods we calculated Bayes factors (Kass and Raftery 1995) to compare CID-3 against the MuSSE-3 model. To approximate the marginal log-likelihood we calculated 18 stepping stones using the methodology from Xie et al. (2011). We calculated the difference between the log-marginal likelihoods of CID-3 and MuSSE-3 models using the statistic *κ* defined as *κ* = *ln*(*P*(*X*|*CID* − 3)) − *ln*(*P*(*X*|*MuSSE* − 3)), where *X* represents our state sample and the phylogenetic tree. If *κ* has a value larger than 1, then the CID-3 model is preferred. The MuSSE-3 model is preferred when *κ* < −1. The test statistic is inconclusive when *κ* has a value in the interval (−1, 1).

### Diversification models with different assumptions about extinction rates

As the fossil record for passerines is relatively sparse (see e.g., Mayr 2005; Tyrberg 2008) there is limited paleontological information from which to estimate extinction rates and species turnover over time. However, some fossil evidence for passerines suggests that extinction rates and turnover have been high. This evidence includes the extinctions of entire early lineages (Hieronymus et al. 2019; Ksepka et al. 2019), high turnover within Europe between the Oligocene and the present (Mayr 2005; Manegold 2008; Bochenski et al. 2021), and Pleistocene-Holocene species extinctions (Rando 1999; Seguí 2001; Claramunt and Rinderknecht 2005; Rando et al. 2010; Oswald and Steadman 2015; Stefanini et al. 2016; Rando et al. 2017; Steadman and Oswald 2020). In addition, evidence from molecular phylogenies of extant taxa suggests passerine turnover has been high (Greenberg et al. 2021). Unfortunately, diversification models fit to phylogenetic trees of extant taxa, as in this study, can greatly underestimate extinction rates (Quental and Marshall 2010; Höhna et al. 2011; Louca and Pennell 2021), producing errors in diversification analyses (Stadler 2013). Thus, we sought to examine whether our results were robust to assumptions about extinction rate. This approach may be useful more generally, as classic paleontological work on extinction rates (Raup 1992; Sepkoski 1998,see also Marshall 2017) has suggested that speciation rates tend to be only slightly larger than extinction rates for clades with net positive diversification rates, and state-dependent diversification models generally have not accounted for this possibility.

In a Bayesian framework, one way to incorporate knowledge about higher extinction rates is by using a prior distribution that accommodates independent information about the magnitude of the rates. Therefore, in our three-state, and hidden-state models, we defined prior distributions for the extinction rates with elevated lower bounds, and we systematically increased these lower bounds across model fits (Table A1). We specified these lower bounds by making extinction rates’ lower bound dependent on speciation rates. Hence, in mathematical form, we defined this process by using the linear function for extinction rates as *μ*_*i*_ = *A* * *λ*_*i*_ + *δ* where *A* = 0.7, 0.8, 0.9, 1, 1.1, for all the states *i* = *D, C, H* (Fig. 2d). We also explored the effect of enhanced extinction in the MuHiSSE-3 model by allowing *A* = 0.8 (Fig. 2e). Defining the extinction rate prior distribution as a linear combination of random variables allowed us to fix the value *A* as the lower bound for the extinction fraction 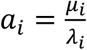 (using use the same definition of extinction fraction as Beaulieu and O’Meara 2016) while permitting extra variability in the diversification rates so that extinction rates are not fully a deterministic function of speciation rates. For example, an extinction rate of dome-nesters that is at least 80% of the speciation rate is defined using the equation *μ*_*D*_ = * *λ*_*D*_ + *δ*. In this example, the posterior estimation of the extinction fraction for state Dome 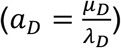 is forced to be at least 80%, but the extinction fraction *a*_*D*_ can still be larger than the lower bound of 0.8 thanks to random variable *δ*, that is 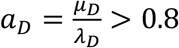. In most of our models, prior distributions for extinction rates were defined equally (i.e. *μ*_*D*_ = *μ*_*C*_ = *μ*_*H*_=*A* * *λ*_*D*_ + *δ*). In one model, we specified *μ*_*H*_ < *μ*_*D*_ < *μ*_*C*_ by making *A* = 0.6, 0.7, and 0.8 respectively to verify that: 1) our inferences weren’t driven by this specification of the prior; and 2) that likelihood is informative given our dataset (supplementary Fig. S14).

Overall, this choice in prior distributions allows us to restrict the parameter space and find estimates of diversification assuming the extinction fraction is high. Linking speciation and extinction rates does not change the dynamics of speciation and extinction in SSE models because speciation and extinction parameters in birth and death models are functionally linked by definition (Kendall 1948; Feller and Feller 1968; Stadler and Bokma 2013). An in-depth discussion on how speciation and extinction parameters in SSE models are functionally linked can be found in Helmstetter et al. (2021).

### Ancestral state reconstruction using diversification-free models

We estimated the most probable state at the root, since previous studies have found that hole-nesting is the most likely the state at the root. The results showed that nest type evolution is not linked to the diversification process (see Results section).

Therefore, we defined a three-state (Mk3, Fig. 2f) Markov model without diversification parameters (hereafter, “diversification-free model” referring to the lack of diversification parameters in the model) to reconstruct the ancestral nest type in passerines. We calculated the marginal ancestral state reconstruction at each of the internal nodes.

Since our reconstructions use a Bayesian framework, in each of the nodes we plot the state with the maximum *a posteriori* value of the marginal posterior distribution (Fig. 5), as previously done in Freyman and Höhna (2018), and Zenil-Ferguson et al. (2019), and implemented in RevBayes (Höhna et al. 2016). For all our models in Figure 2, we assume the root values are stochastic, and are modeled using a Dirichlet probability distribution with frequency parameter (1/3,1/3,1/3) as the prior distribution (see the number of parameters in Fig. 2). We calculated the posterior distribution for this stochastic vector via our MCMC algorithms.

### Implementation of models and inferences

All the diversification and diversification-free models in Fig. 2 were implemented as graphical models in RevBayes software (Höhna et al. 2016). We customized and ran a Markov chain Monte Carlo (MCMC; (Metropolis et al. 1953; Hastings 1970) algorithm to sample the posterior distribution of each model. For all the models in Figure 2, we assumed that the state value at the root was unknown, and we estimated the posterior distribution of the frequencies at the root. Convergence and effective sample sizes of at least 250 for every parameter in the MCMC were assessed using Tracer (Rambaut et al. 2018) and two chains were run per model to verify convergence. Each MCMC run was performed in the HiPerGator cluster at the University of Florida, and took an average of 240 hours to converge. All our implementations are available at https://github.com/roszenil/nestdivmodel (will be updated to Dryad for SysBio).

## Results

### Diversification is not linked to nest type

While our MuSSE-3 model found associations between nest type and diversification rates, our MuHiSSE-3 model showed that these associations were spurious (Fig. 3). This is because the addition of hidden states indicates that diversification is driven by unmeasured factors other than nest type. In particular, for the MuSSE-3 model of nest evolution, we found that hole nesters had faster or equal net diversification (defined as speciation minus extinction) than cup nesters, and the smallest net diversification was associated with dome nesters (Fig. 3a). This result was also true when the prior of extinction fraction was faster (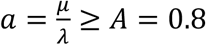, Fig. 3a), and for all elevated extinction fractions (0 < *A* ≤ *a* = *μ*/*λ* < 1) (Fig. 2d is the model and estimation is in Fig. A1). When fitting the hidden state model MuHiSSE-3 (Fig. 3b), we found that the three-state posterior distributions of net diversification rates completely overlap within hidden states A and B while being different between A and B, indicating that the diversification rate differences across the phylogenetic tree are due to hidden states (Fig. 3b). These results were consistent when we allowed MuHiSSE-3 to have an elevated extinction fraction of at least 0.8 (Fig. 2e shows model and estimation in Fig. 3b). Furthermore, when calculating Bayes factors comparing the CID-3 (diversification rates independent of nest type, or null hypothesis) model against the MuSSE-3 (diversification rates depend on nest type), we found that the CID-3 model is preferred over the MuSSE-3 (*κ* = −6033.72 − (−7772.40) = 1738.67 > 1). Altogether, there is strong evidence that the heterogeneity observed in passerine diversification rates is not linked to nest type.

**Figure 3.**
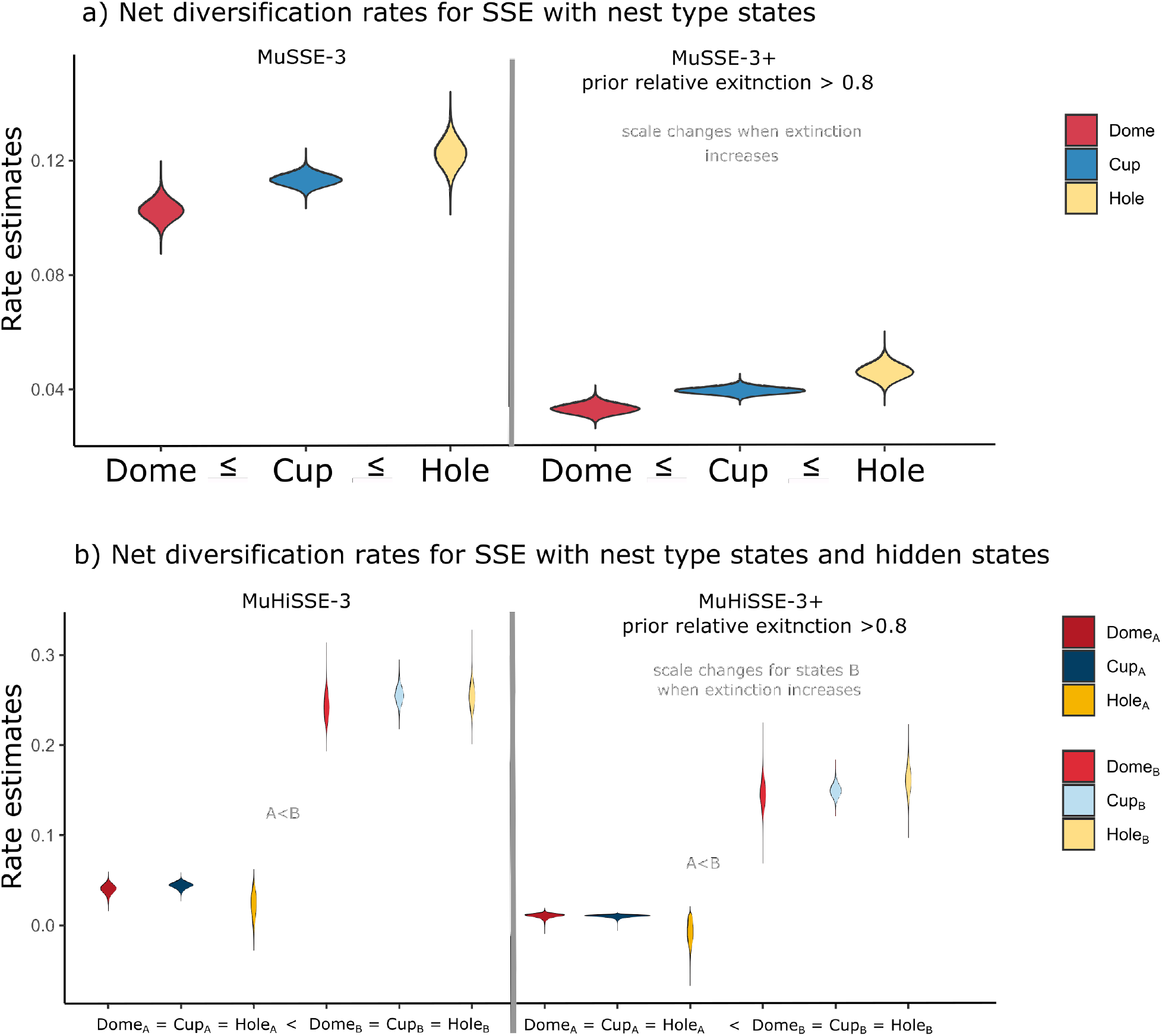
Net diversification rate estimates (speciation and extinction) for a) multistate dependent diversification models (MuSSE-3), and b) multistate and hidden state dependent diversification models (MuHiSSE-3). In models in (a) we consistently estimated that hole nesters have faster net diversification rates than dome nesters, and that cup nesters had intermediate rate estimates. This finding was consistent when we assumed the extinction fraction to be at least 0.8 (MuSSE-3 model + prior extinction fraction > 0.8). The most noticeable change is that increasing the extinction fraction (MuSSE-3 model+ extinction fraction > 0.8) decreases net diversification estimates (y-axis), but the relative differences between states are maintained. For models in (b) we observe that all diversification differences are due to the differences in hidden state (A<B) rather than nest type (Dome=Cup=Hole for A and B states), which is strong evidence against the direct influence of nest type on the speciation and extinction of passerines. Even increasing the extinction fraction (MuHiSSE-3 + prior extinction fraction > 0.8) yields the same results.

### Evolving out of holes is faster than evolving back into them

Since nest type is not linked to the diversification process, we assessed the dynamics under the diversification-free Mk3 model. The posterior distribution of transition rates from the Mk3 model shows that the rate of transitioning from hole to cup was the fastest (mean *q*_32_ = 9.1×10^−3^, 95% credible interval (6.5×10^−3^, 0.01), followed by the transition from hole to dome (mean *q*_31_ = 6.5×10^−3^, 95% credible interval (3.7×10^−3^, 8.2×10^−3^), Fig. 4). Transition rates from either dome or cup to hole nests were similar, and both were slower than the transition rates out of hole nests (dome to hole transition rates have a mean *q*_13_ = 1.6×10^−3^, and 95% credible interval (6.6×10^−4^, 2.8×10^−3^); cup to hole transition rates have a mean *q*_.3_ = 1.2×10^−3^, credible interval (7.3×10^−4^, 2.5×10^−3^), shown in Fig. 4.

**Figure 4.**
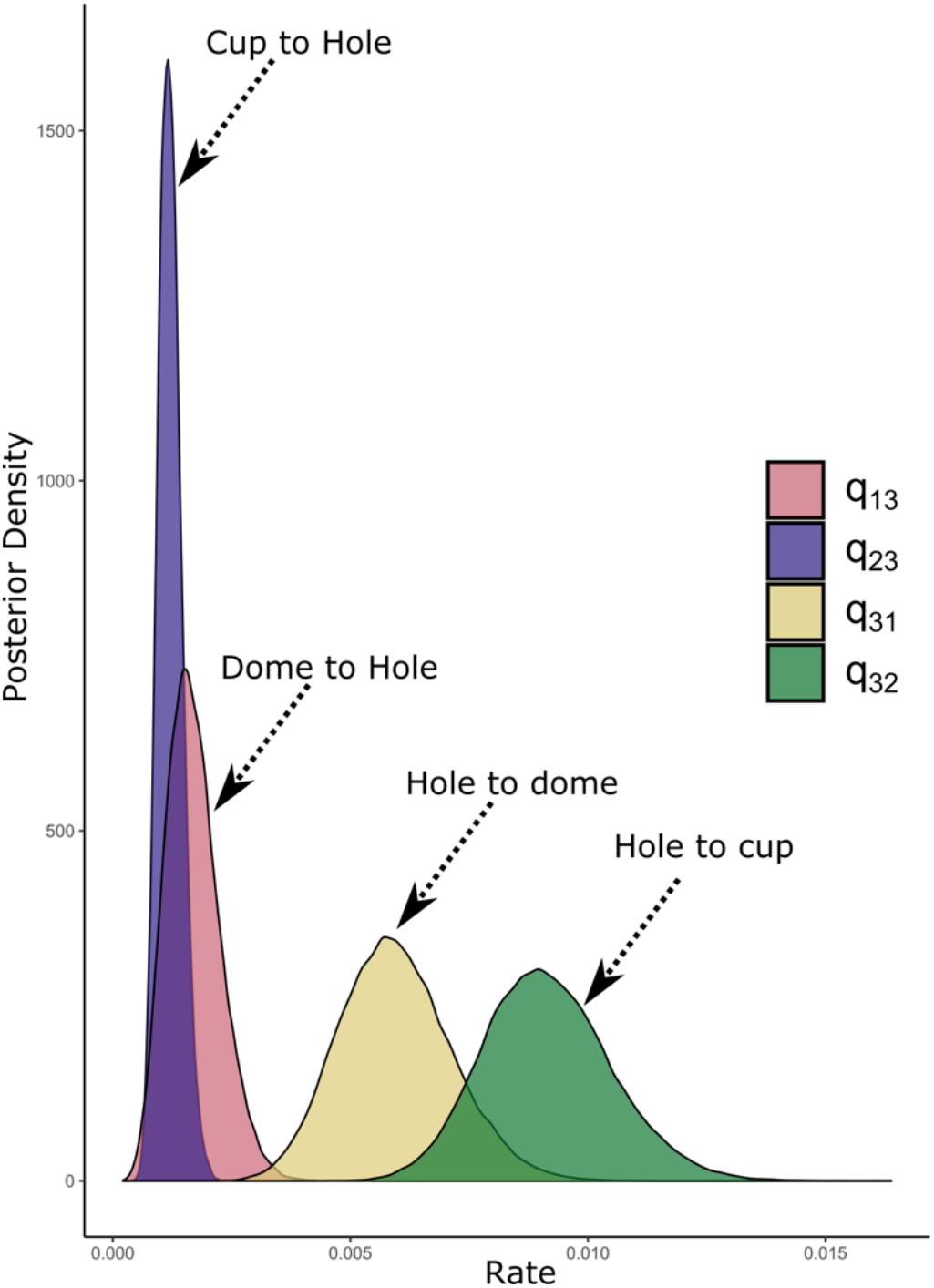
Transition rates between nest states for diversification-free models in the model with three states (Mk3). The posterior distribution for the rate from hole to cup (*q*_*3*2_) is faster than any other rate to and from hole. We found that evolving out of hole is faster than into hole, with hole to cup faster than hole to dome.

### Hole-nesting is the most probable state at the root is hole when including outgroup

The Mk3 (Fig. 2f) model reconstructs hole-nesting as the state with the highest probability at the most recent common ancestor of the Passeriformes. We found that the maximum *a posteriori* for the root’s marginal posterior probability was 0.77 for hole-nesting when the tree included the parrot *Strigops habroptila* as the outgroup, and 0.7 for the node that ancestor of the passerines only. This probability decreased to 0.41 when the tree only contained passerines and no outgroup (states dome and cup make the other 0.51 of probability, Fig. S13 in supplement). The multivariate posterior distribution for the stochastic vector at the root had as its maximum the vector (Dome=0.30, Cup=0.25, Hole=0.45; Figure 5).

**Figure 5.**
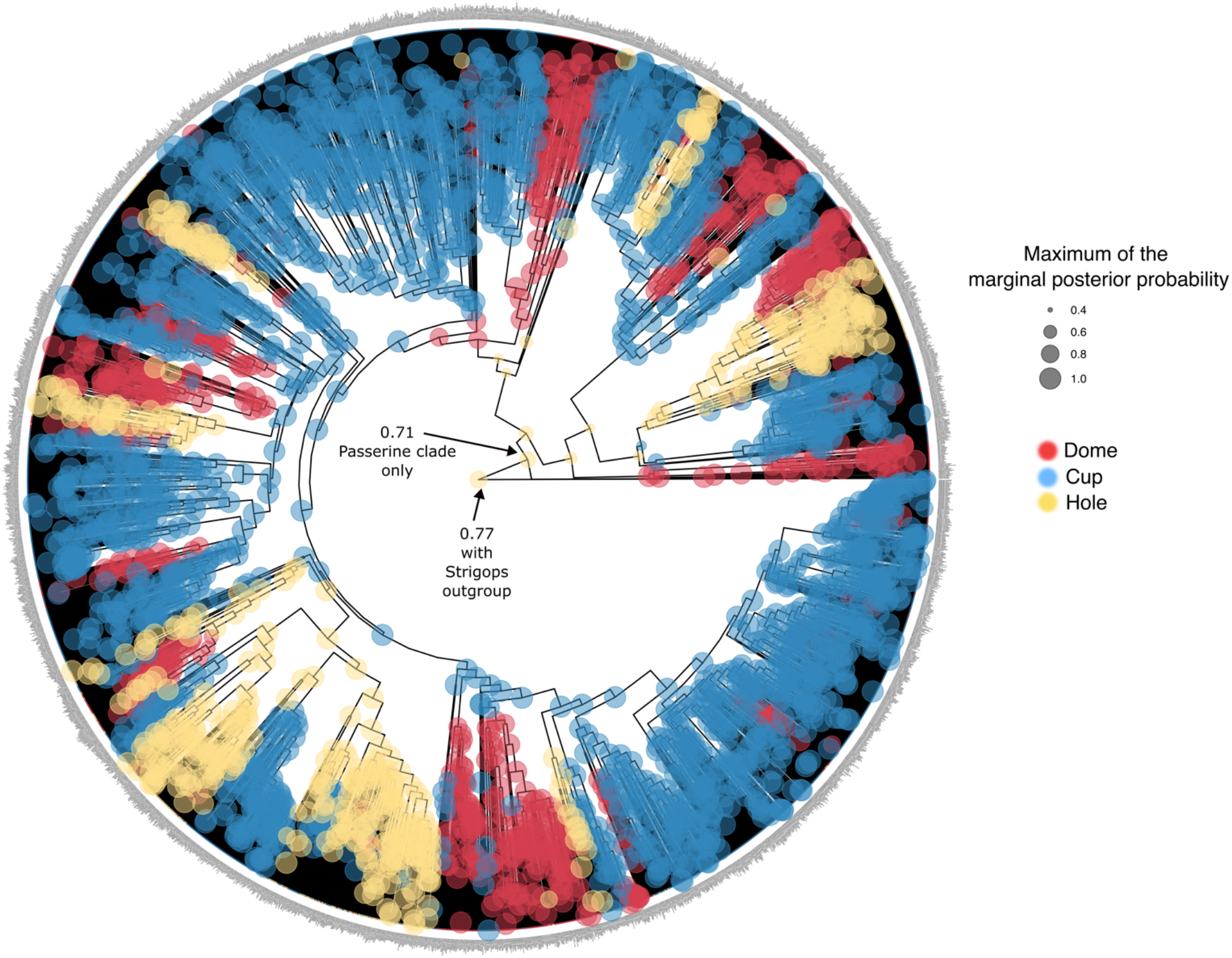
Ancestral state reconstruction for nest type using a Markov model with three states (Mk3). The size of a node represents the maximum probability value of the marginal posterior distribution of the node. The color of the node represents the nest type associated with the maximum posterior probability. This tree includes the hole-nesting *Strigops habroptila* as the outgroup. Figure made using RevGadgets (Tribble et al. 2022).

## Discussion

Over the years, there have been diverse perspectives on the link between ecological specialization and evolutionary diversification (Vamosi et al. 2014). On the one hand, ecological specialization has been viewed as an evolutionary dead end because it leads to evolutionary changes in traits that are difficult to reverse, which can leave specialized taxa at higher risk of extinction when conditions change. On the other hand, other studies have emphasized the links between specialization and increased diversification (Losos et al. 1994; Schluter 2000) because a narrower niche could increase diversification either directly, if niche shifts are associated with multiple speciation events (Yoder et al. 2010, Velasco et al. 2016), or indirectly, because of differences in dispersal, connectivity, population persistence, and/or range size of specialist and generalist species (Greenberg and Mooers 2017; Harvey et al. 2019). Testing the outcomes of specialization on diversification rates has been difficult because macroevolutionary models require datasets with a large number of independent origins of specialization (Day et al. 2016).

Here, using state-dependent diversification models in a large tree of passerines, we found that hole-nesting specialization does not differ from cup and dome-nesting in diversification rates. We also found that transition rates into hole-nesting were low compared to transitions out of hole-nesting (Fig. 4), and that the most probable state for the root of all passerines was hole-nesting. These three results highlight the lack of support for the hypothesis that specialization leads to an evolutionary dead end. Our results, combined with a number of recent studies addressing this hypothesis (Day et al. 2016; Cyriac and Kodandaramaiah 2018), raise the question of whether the link between diversification rates and ecological specialization is truly variable across taxa and traits, or whether newer phylogenetic comparative methods and larger data sets are allowing us to test hypotheses more rigorously. Interestingly, in our study, which uses a large dataset with numerous transitions to the specialized hole-nesting state, we found that, in the absence of accounting for hidden states, adoption of hole-nesting behavior led to higher, not lower, diversification rates (Fig. 3a). However, when we included hidden states, and thus accounted for the possibility that other, unmeasured variables are driving the relationship, we found no difference in diversification rates across the three nesting types (Fig. 3b). This comparison emphasizes that phylogenetic studies of specialization that do not account for hidden states may need to reevaluate their null hypotheses of diversification. For example, a recent study by Medina et al. (2022) concluded that there are differences in extinction rates between cup and dome nesters, but hidden state models in this study were not presented due to failed convergence, and extinction rates cannot be interpreted independently from speciation rates (Helmstetter et al. 2021), so the link between nest type and diversification remained untested. Our results, in combination with recent tests of the evolutionary dead end hypothesis, show that there is growing evidence that specialization rarely leads to an evolutionary dead end (Stern et al. 2017; Villastrigo et al. 2020).

### Modeling the diversification consequences of specialization

State-dependent diversification models are a flexible tool to test whether specialization is linked to diversification rates. A recent ongoing discussion in the field of macroevolution centers on the impossibility of estimating speciation and extinction rates from phylogenetic trees with only extant taxa using birth and death stochastic models with time-dependent parameters (Louca and Pennell 2020). The non-identifiability of parameters in the time-dependent diversification models, and the possibility of multiple congruent likelihoods across time-dependent models may affect the state-dependent diversification models presented here. State-dependent diversification models are members of the class of the time-dependent diversification models (with time being a constant function) explored in Louca and Pennell (2020). However, taking into consideration the nest type, and using informative priors to represent different extinction scenarios (proposing other constant functions for rates at higher levels), shows the potential for recovering the direction of differences between states, even if point estimates for speciation and extinction rates are not the same across these scenarios (Fig. 4, and Fig. A1), as suggested by Louca and Pennell (2020). Our main findings under the assumptions of enhanced extinction suggest that the relative differences among states are possible to infer despite the non-identifiability between different extinction histories. Therefore, it is important when applying state-dependent diversification models to clarify whether the goal is to obtain point estimates for speciation or extinction rates or to find relative differences in the history of diversification linked to the states. In this study, we were interested in the latter, and our results suggest that finding relative differences of diversification between states can be performed in a systematic fashion with consistent results.

Overall, we were able to rigorously test different hypotheses and extinction scenarios because of the size of the tree, the number of transitions between nest types, and the large proportion of passerines in the sample (> 51%). Studies of specialization linked to diversification might find spurious associations when independent transitions to and from the specialized state are too few (Uyeda et al. 2018), tree size is not large enough (Davis et al. 2013), sampling fraction is small (Chang et al. 2020), or when hidden state models are not incorporated to test for the significance of trait-dependent diversification (Beaulieu and O’Meara 2016; Caetano et al. 2018). Historically, standards for sample quality and interpretation of the parameters as a joint function of speciation and extinction in SSE models have been lacking (Helmstetter et al. 2021), so it is important that future studies of state-dependent diversification be aware of these issues.

### Ecological and evolutionary consequences of specialization

The ecological consequences of specialization can be diverse – specialization can, in some cases, decrease competition for resources and, in other cases, increase it. For example, many studies have focused on diet or host specialization, which frequently reduce competition over food or breeding resources (Vamosi et al. 2014). However, specialization can also enable species to escape predation (e.g., Singer et al. 2019) and, in such cases, may actually increase competition for scarce, protective resources. Such is the case for hole-nesting birds, where an important consequence of hole-nesting is that it strongly reduces the risk of nest predation, an important driving force behind life history evolution in birds (Martin 1995). In our study, we were able to distinguish between the influence of ecological specialization per se versus one specific ecological consequence of specialization – reduced predation. Dome-nesting birds also show decreased predation rates (Oniki 1979b; Linder and Bollinger 1995; Auer et al. 2007; Martin et al. 2017), but they are relatively unspecialized in nesting types compared to hole-nesters. However, neither hole-nesting nor dome-nesting were associated with elevated diversification rates. This suggests that, in passerine birds, reduced nest predation rates do not have a strong influence on diversification dynamics and, while escaping predation can often lead to ecological release (Herrmann et al. 2021), this does not necessarily translate into increased ecological opportunity or subsequent adaptive diversification. Moreover, reduced predation can also increase species’ ability to persist and, hence, decrease extinction risk. Yet, a change in abundance of nesting holes, such as declines in tree cavities, has been shown to lead to heightened risk of extinction (Duckworth and Badyaev 2007). Thus, the lack of a relationship with diversification rates may reflect a balance between these various ecological consequences of adopting hole-nesting.

Another factor that may account for inconsistency across studies in the links between specialization and macroevolutionary dynamics is the extent to which specialization results in further evolution of specific traits. Ecological specialization is a narrowing of an organism’s niche, and so results from a change in how an organism interacts with its environment. Futuyma and Moreno (1988) point out that specialization, at a minimum, only requires a behavioral shift, and specialized taxa vary in the extent that there is subsequent evolution of other traits. Thus, it is possible that whether specialization is an evolutionary dead end or not may be strongly linked to the extent of secondary adaptation that follows it. In the case of hole-nesting birds, specialization is largely a behavioral shift, although it does lead to evolutionary changes in many life history and breeding traits (see below). Thus, it may be that hole-nesting specialization does not lead to an evolutionary dead end because of the relative ease of reversing these traits (Fig. 4). The low rate of evolutionary transitions into, relative to out of, hole-nesting (Fig. 4) provides further evidence against the evolutionary dead end hypothesis and instead suggests that evolving ecological specialization can be challenging. In the case of hole-nesters, becoming ecologically specialized involves facing increased interspecific competition, as holes are sought not only by other birds but also by other vertebrates (Newton 1994). Thus, competition may prevent evolutionary transitions towards specialization. Evolutionary transitions from hole-nesting to open-cup nesting, which occur at a relatively high rate, involve different evolutionary challenges, such as life history specialization. Open-cup nesting species have exceptionally high nestling growth rates, which appear to be an adaptation to increased predation risk (Ricklefs 1979). Moving from the relaxed selection regime on growth rates associated with hole-nesting to the strong selection regime on growth rates in open-cup nests would seem to act as a filter that would limit this transition. However, our results show that, at least in passerine birds, evolving such life history shifts is easier than evolving the ability to deal with the intense competition of hole-nesting.

We also found evidence that it was easier for birds to transition from hole to cup rather than hole to dome. Most hole nesters build an open-cup style nest, rather than a dome, within the cavity (Price and Griffith 2017). Our finding that transitions to open-cup nesting were easier compared to transitions to dome may simply reflect that transitions from hole-nesting more often occurred in taxa that were already building open-cup nests within their holes. Thus, for most hole nesters, transitioning to dome-nesting may be more difficult because it would require two steps: leaving cavities and changing how the nest is built. While in the minority, there are several clades, particularly at the base of the passerine tree, that build dome style nests within cavities (Price and Griffith 2017). Future studies comparing transitions to dome and open-cup nests based on whether hole-nesting species are already building a dome or cup style nest within their cavity would add further insight into the mechanisms behind these different transition rates.

### Ancestral State Reconstruction

In our ancestral state reconstruction, we found that hole-nesting is the most probable state at the root of the passerines when including the parrot *Strigops habroptilus* as the outgroup. Using this phylogeny, we found that the maximum of the marginal posterior distribution of the root node had a probability of 0.77 for hole-nesting, and the root node of the passerine clade had a 0.7 probability of being hole-nesting (Fig. 5). Without this outgroup, we still recovered hole-nesting with a maximum a posteriori of 0.41 (see Fig. S13 in supplementary information) but probability 0.51 for dome nesting. Previous work has emphasized that hole-nesting, dome-nesting, and open-cup nesting all appear to have occurred early in the history of passerines (Collias 1997; Price and Griffith 2017; Fang et al. 2018; McEntee et al. 2018), with Collias (1997) suggesting that the nest type of the earliest passerine might be unknowable because of the apparent rapid evolution of nest type in early passerines. Our ancestral state reconstruction, using an approach where we included strong species-level sampling and assessed whether differing diversification rates had to be accounted for (Maddison 2006) tips the balance slightly in favor of hole-nesting as the ancestral state for the common ancestor of extant passerines when using nest type information for outgroups. Notably, this result contrasts with the ancestral state reconstruction of Fang et al. (2018), which included more sampling outside of passerines and less species-level sampling within passerines. Hole-nesting as the root state for passerines might seem counterintuitive if the definition of specialization is confounded with the concept of a “derived state”. In the original definition of specialization by Futuyma and Moreno (1988) it is stated that specialization is based on ecology and function in an ecosystem; therefore, we shouldn’t expect that a specialized state is a derived one. In our study, support for hole-nesting as the ancestral state serves to underscore how radically the evolution of hole-nesting as a specialization fails to meet the generalists-to-specialists view of evolution; rather than a dead end, all passerine diversity emerged from a specialist ancestor. When we excluded the *Strigops* outgroup we found that the most probable state at the root is dome-nesting (Fig. S13). This second result shows that there is significant evidence against cup-nesting for passerine and that ancestral state reconstructions are heavily influenced by the inclusion of exclusion of lineages. In the future, more sampling of passerine lineages and their nest types will have a powerful effect in resolving the root state.

## Conclusions

Our macroevolutionary analysis failed to find any link between specialization of nest type and diversification rates under different extinction scenarios, suggesting that there is little support for the evolutionary dead end hypothesis in this case study.Moreover, contrary to this hypothesis, we also found that transitions from the specialized state were relatively easy compared to transitions toward the more generalist states and that the root state of passerines is most probably hole-nesting. Our results suggest that the ecological consequences of resource specialization,whether due to escape from competition or predation, might be key to understanding its macroevolutionary consequences. This work adds to other recent studies that have found little support for the evolutionary dead end idea, suggesting that evolution of resource specialization is more evolutionarily labile than previously thought. We suggest that future studies of this question would benefit from explicit comparison of resource specialization that vary in their ecological consequences, as well as the extent of trait evolution necessary for specialization. Such studies would enable a greater understanding of the mechanisms that underlie variable links between specialization and macroevolutionary dynamics.

## Acknowledgements

We would like to thank A.E. Sara Ruane and two anonymous expert reviewers for their insightful comments. We are grateful for Dr. Sonal Singhal who offered feedback on an initial draft of this work, Dr. Carrie Tribble who helped with RevGadgets visualizations, and Jenna McCullough for the illustration of nests. This work started and concluded during the toughest time of the COVID-19 emergency, slow progress was made during two years thanks to the support of many of our colleagues and families. RZF is supported by NSF-DEB 2055466.

## Appendix

**Table A1.** Assumptions of extinction in different MuSSE-3 models.

| **Table A1.** Assumptions of extinction in different MuSSE-3 models. |  |  |  |
| --- | --- | --- | --- |
| **Assumption**<br>**Extinction fraction** | **Extinction as a**<br>**function of**<br>**speciation** | **Prior distributions**<br>**used\*** | **Interpretation** |
| 1. NA<br>(Fig. 2a) | NA | $\lambda_i \sim \text{LogN}(\mu_\lambda, \sigma_\lambda)$<br>$\mu_i \sim \text{LogN}(\mu_\mu, \sigma_\mu)$ | Speciation and extinction freely estimated |
| 2. $\mu_i/\lambda_i \geq 0.7$<br>with $i = C, D, H$<br>(Fig. 2d) | $\mu_i = 0.7 \lambda_i + \delta$ | $\lambda_i \sim \text{LogN}(\mu_\lambda, \sigma_\lambda)$<br>$\delta \sim \text{Exp}(2)$<br>$i = C, D, H$ | Extinction rate is at least 0.7 of speciation rate |
| 3. $\mu_i/\lambda_i \geq 0.8$<br>with $i = C, D, H$<br>(Fig. 2d) | $\mu_i = 0.8 \lambda_i + \delta$ | $\lambda_i \sim \text{LogN}(\mu_\lambda, \sigma_\lambda)$<br>$\delta \sim \text{Exp}(2)$<br>$i = C, D, H$ | Extinction rate is at least 0.8 of speciation rate |
| 4. $\mu_i/\lambda_i \geq 0.9$<br>with $i = C, D, H$<br>(Fig. 2d) | $\mu_i = 0.9 \lambda_i + \delta$ | $\lambda_i \sim \text{LogN}(\mu_\lambda, \sigma_\lambda)$<br>$\delta \sim \text{Exp}(2)$<br>$i = C, D, H$ | Extinction rate is at least 0.9 of speciation rate |
| 5. $\mu_i/\lambda_i \geq 1.0$<br>with $i = C, D, H$<br>(Fig. 2d) | $\mu_i = \lambda_i + \delta$ | $\lambda_i \sim \text{LogN}(\mu_\lambda, \sigma_\lambda)$<br>$\delta \sim \text{Exp}(2)$<br>$i = C, D, H$ | Extinction rate is greater or equal to speciation rate |
| 6. $\mu_i/\lambda_i \geq 1.1$<br>with $i = C, D, H$<br>(Fig. 2d) | $\mu_i = 1.1 \lambda_i + \delta$ | $\lambda_i \sim \text{LogN}(\mu_\lambda, \sigma_\lambda)$<br>$\delta \sim \text{Exp}(2)$<br>$i = C, D, H$ | Extinction rate is at least 1.1 of speciation rate |
| 7. $\mu_C/\lambda_C \geq 0.8$<br>$\mu_D/\lambda_D \geq 0.7$<br>$\mu_H/\lambda_H \geq 0.6$<br>(Results in supplementary information Fig. S12) | $\mu_C = 0.8 \lambda_C + \delta$<br>$\mu_D = 0.7 \lambda_D + \delta$<br>$\mu_H = 0.6 \lambda_H + \delta$ | $\lambda_i \sim \text{LogN}(\mu_\lambda, \sigma_\lambda)$<br>$\delta \sim \text{Exp}(2)$<br>$i = C, D, H$ | For cup, extinction rate is at least 0.8, for dome, is at least 0.7, and for hole, extinction rate is at least 0.6 of speciation rate. |
| *Prior distributions for log-speciation were specified as Normal distributions with parameters $\left(\eta_\lambda = \ln\left(\ln\left(\frac{\ln(3225/2)}{\text{Tree Age}}\right)\right), \nu_\lambda = 1.12\right)$ in RevBayes. This implies that speciation parameters are distributed log-Normal $\lambda_i \sim \text{LogN}(\mu_\lambda = 0.11, \sigma_\lambda = 3.3)$ (approximate parameters). For parameter ($\delta$) we chose an exponential with rate 2. Other distributions were tested for $\delta$ and results were unaltered. |  |  |  |

In Figure A1 we summarize the results of assumptions 1-6 from table A1. Here we can observe that when it is assumed that the extinction fraction is greater than 0.7 but less than 1, we recover the differences between states (Dome ≤ Cup ≤ Hole). When the extinction fraction is at least 1 and the extinction rate is expected to be about as large as speciation, we found no differences in the posterior distributions of the three states. Finally, when the extinction rate is assumed to be larger than the speciation rate we obtain the reverse effect (Hole ≤ Cup ≤ Dome). These results show that despite our prior assumptions about extinction rate, the hole-nesting taxa will have a greater influence on the direction of net diversification than cup or dome nesters. And finally, the state dome always has the smallest effect on net diversification in absolute value. These results might only be applicable to a trait that does not influence diversification, as we have shown using hidden state and character independent models for nest type (Figure 2). Whether this pattern of finding similar differences in diversification is sustained in the face of enhanced extinction holds for all traits in SSE models remains to be tested.

**Figure A1.**
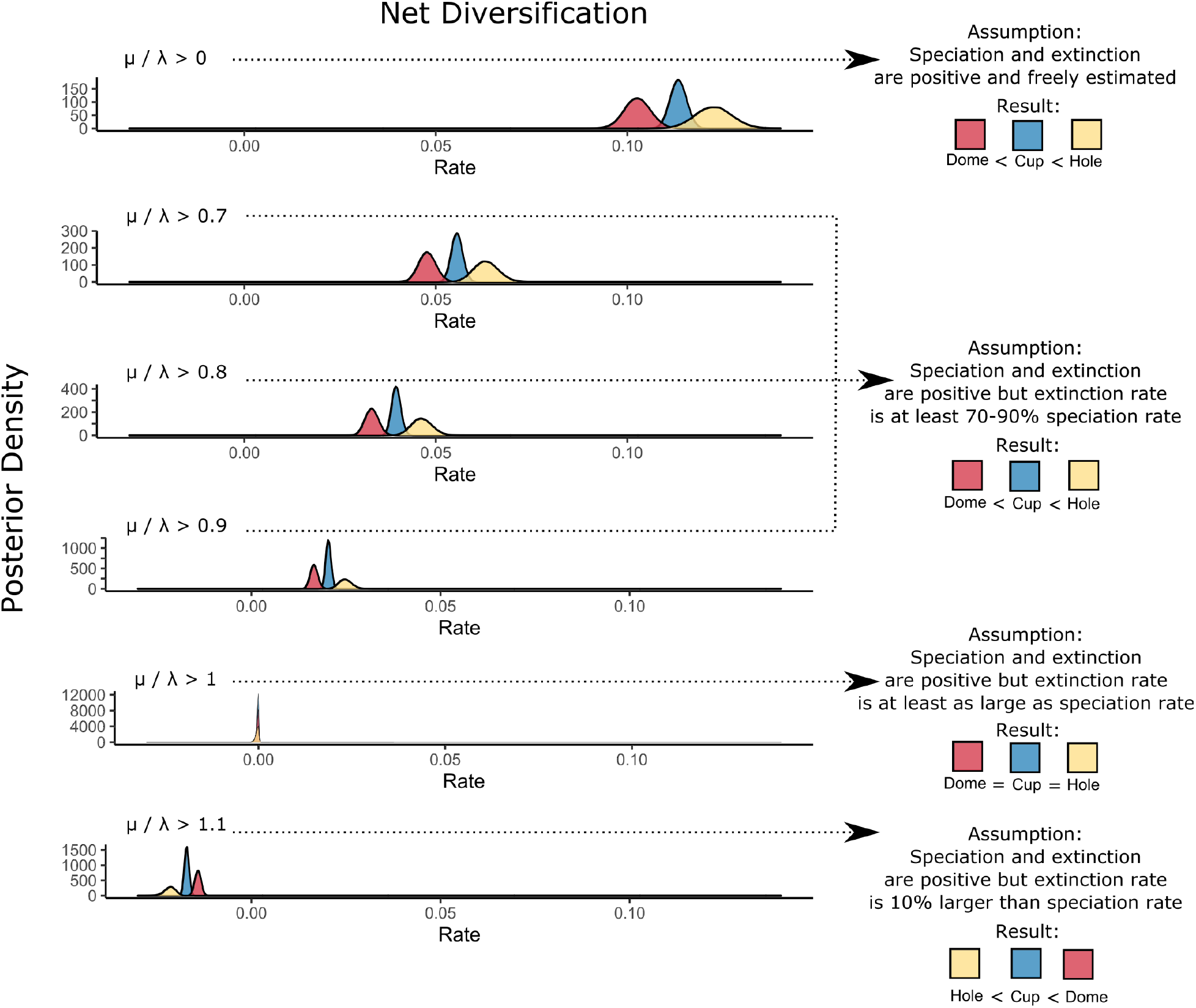
Different *a priori* assumptions about the minimum of extinction fraction 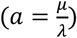. When speciation and extinction are estimated freely (*a* > 0), the posterior distributions for net diversification show that the rate of diversification for hole is faster or equal than cup, and both are faster or equal than dome. This inference is supported even when we assume that extinction rate should be faster (in other words for cases where we fixed a lower bound *A* for extinction fraction (i.e., *a*_*i*_ = *μ*_*i*_/*λ*_*i*_ ≥ *A* = 0.7,0.8, 0.9 with *μ*_*i*_ = *A* × *λ*_*i*_ + *δ* for all states *i* = *D, C, H*) but still smaller than speciation. However, this assumption changes the overall scale leading to smaller net diversification values. The inference from net diversification starts changing when we assumed *a*_*i*_ > 1, that is, that extinction is at least as fast as speciation for every state, where we found that the posterior distributions of net diversifications for every state overlap. Once we assumed that extinction is much faster than speciation (*A* = 1.1), we found a mirrored trend in the inference with hole net diversification being the slowest, and dome being the fastest.

