## Supplemental information for "Linking ecological specialization to its macroevolutionary consequences: An example with passerine nest type"

### Supplementary information for “Linking ecological specialization to its macroevolutionary consequences: An example with passerine nest type”

#### Three state-dependent speciation and extinction models

Complete results for posterior distributions of the different models proposed in Fig. 2a and 2d in the main article shown here.

##### *Full inference for MuSSE-3 model*

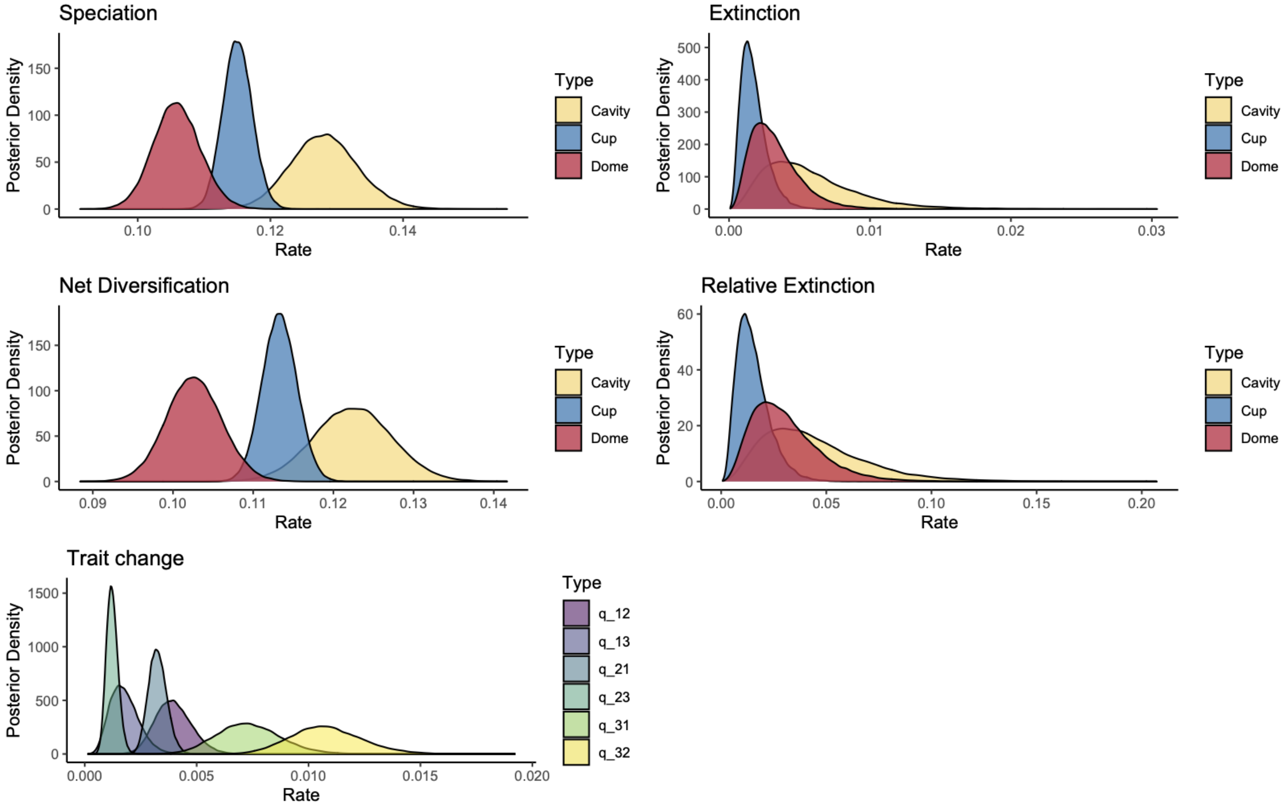


**Figure S1**. Posterior distributions for parameters in MuSSE-3 model. In the main manuscript, the net diversification panel is represented as a violin plot in Fig. 3a (left panel).

***Full inference for MuSSE-3 model  plus prior relative extinction larger or equal than 0.8***

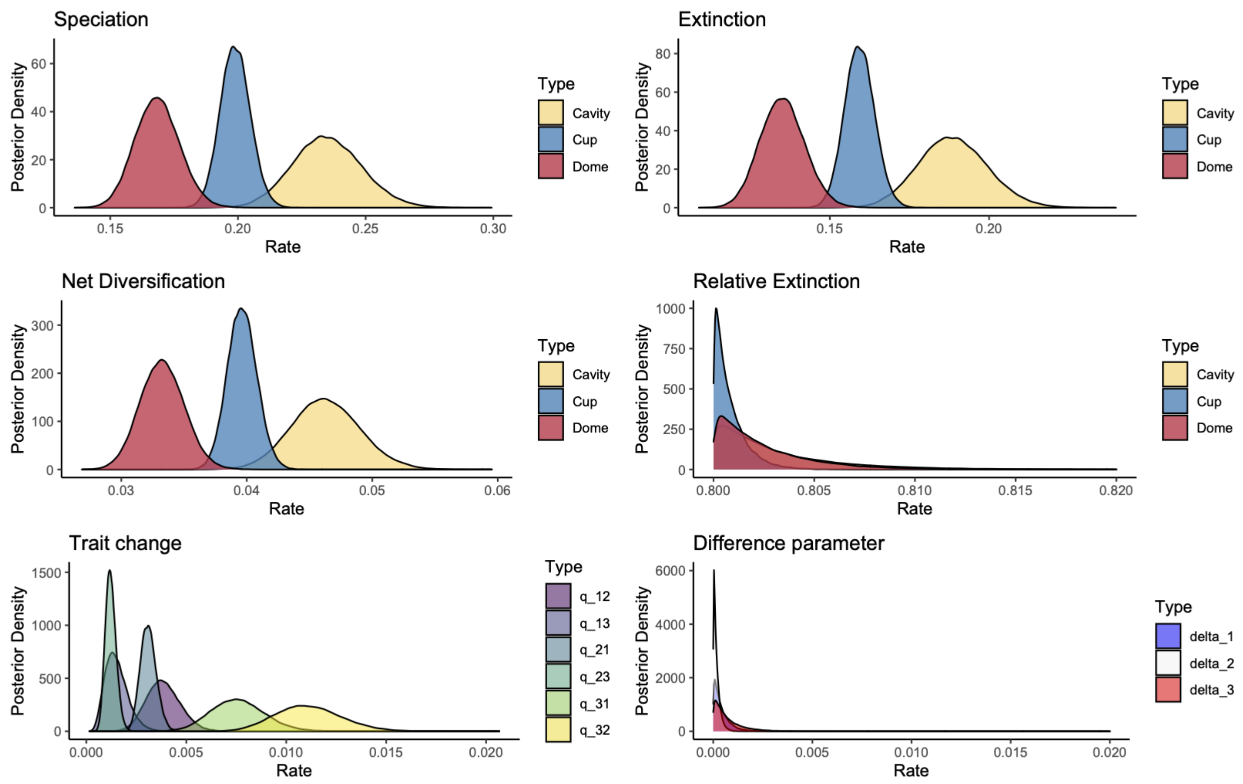


**Figure S2.** The posterior distributions for parameters in MuSSE-3+ prior relative extinction are larger than 0.8. In the main manuscript, the net diversification panel is represented as a violin plot in Fig. 3a (right panel). In this model, the relative extinction rate is assumed to have a minimum value of 0.8 which can be seen on the x-axis of the relative extinction panel

Complete results for posterior distributions of the different models proposed in Fig. 2b and 2e in the main article shown here.

#### ***Full inference for MuHiSSE-3 model***

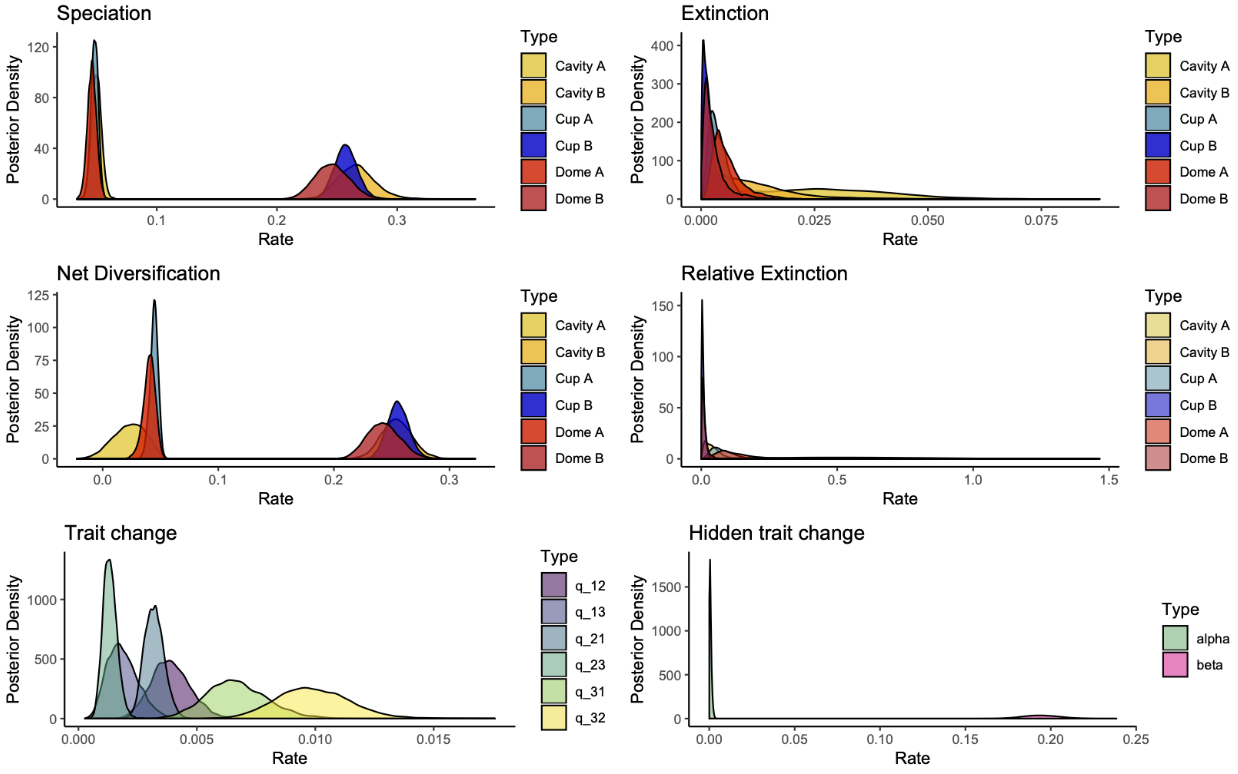


**Figure S3**. The posterior probabilities for parameters of MuHiSSE-3 model. The net diversification rates represented as violin plots in Fig. 3b (left panel) in the main manuscript show that there are no diversification differences among the three states.

***Full inference for MuHiSSE-3 model  plus prior relative extinction larger or equal than 0.8***

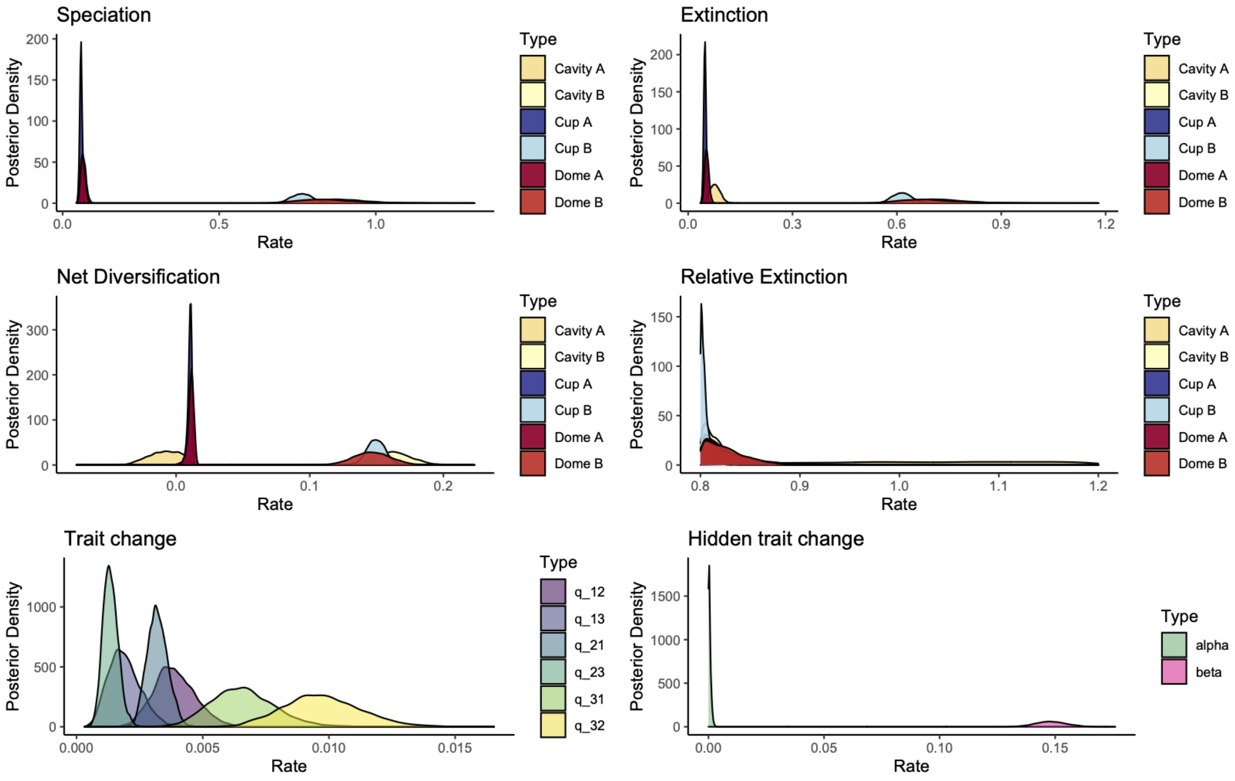


**Figure S4.** Posterior distributions for parameters in MuHiSSE-3 + prior relative extinction larger than 0.8. In the main manuscript, the net diversification panel is represented as a violin plot in Fig. 3b (right panel) resulting in no differences in diversification. In this model, the relative extinction rate is assumed to have a minimum value of 0.8 which can be seen in the x-axis of the relative extinction panel.

**Six state-dependent speciation and extinction models**

The second set of models fitted were six state SSE. MuSSE-6 is a model in which the six main states represent not only dome, cup, and hole nest sites, but also all the pairwise combinations of multiple nest types (dome and cup, dome and hole, and cup and hole , Fig. S5 a) to account for those species that show flexibility in nest type use. MuSSE-6 includes speciation and extinction rate parameters linked only to the the single nest states. Transition rates between all states are possible except transitions between those states representing two different nest types and the single nest type state not including either of these nest types (e.g., state $DC$ does not connect with state $H$ since that represents a double transition, Fig. S5 a). The MuSSE-6 model helped us identify the most appropriate way to score states when taxa can use more than one nest type. Model MuHiSSE-6 (Fig. S5 b) is the extension that includes hidden states, and model Mk6, is a diversification free model (Fig. S5 c).

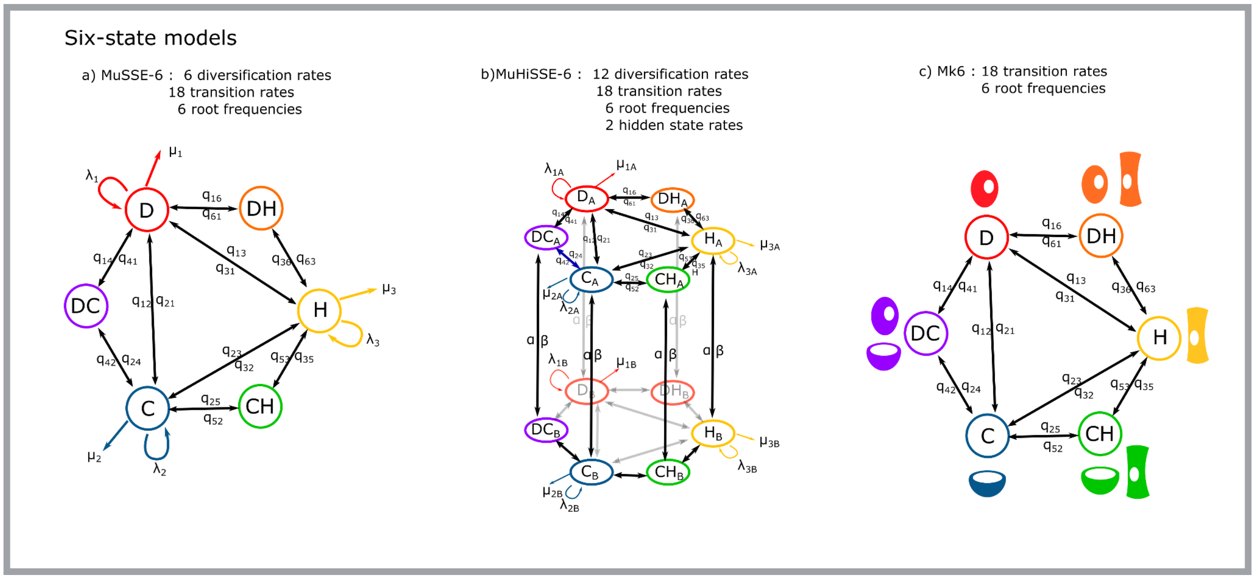


**Figure S5**. Six state speciation and extinction models (a) MuSSE and b) MuHisse- with hidden states), and markov model MK6 (diversification-free).

***Full inference for MuSSE-6 model***

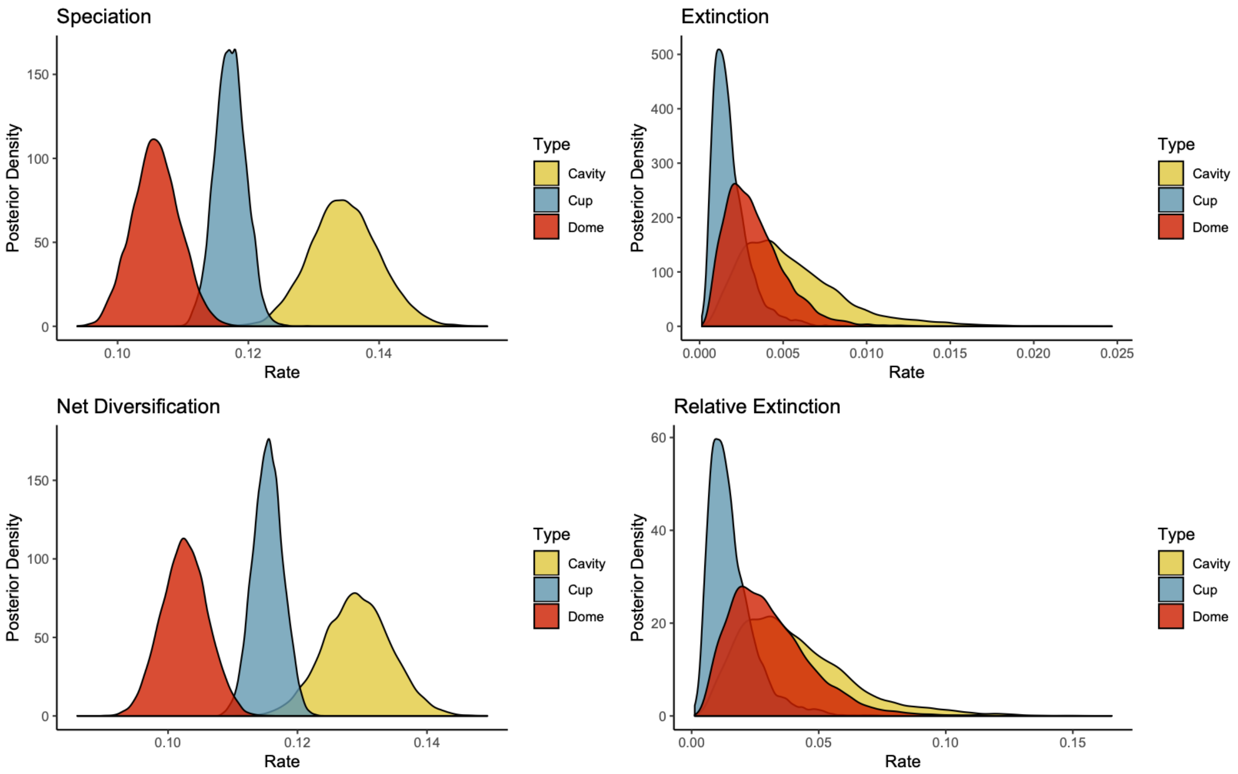


**Figure S6**. Posterior distribution for diversification parameters of MuSSE-6 model. The net diversification panel is represented as a violin plot in the main manuscript in figure 3a.

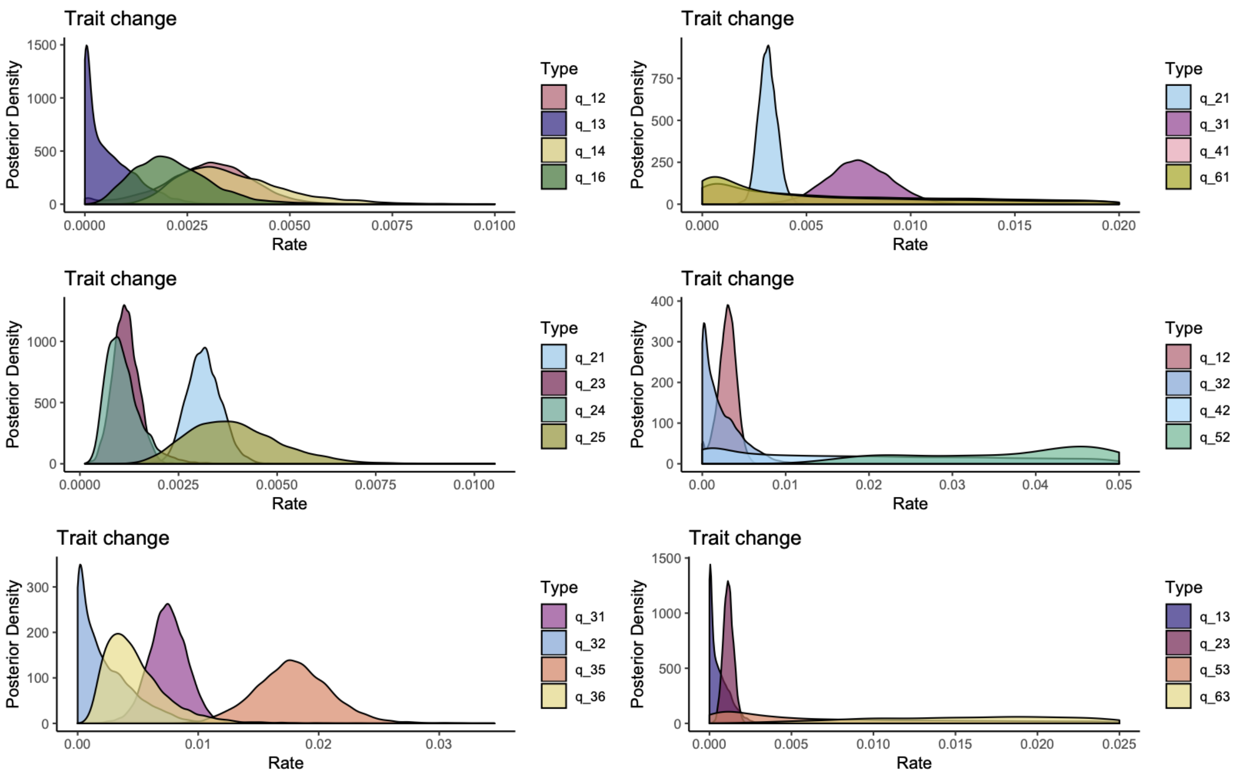


**Figure S7**. Posterior distributions of trait change for model MuSSE-6. The three panels on the first column show the transition rates out of the main three states (dome, cup, and hole respectively) and the three panels on the second column indicate the transition rates towards the main states (dome, cup, and cavity respectively).

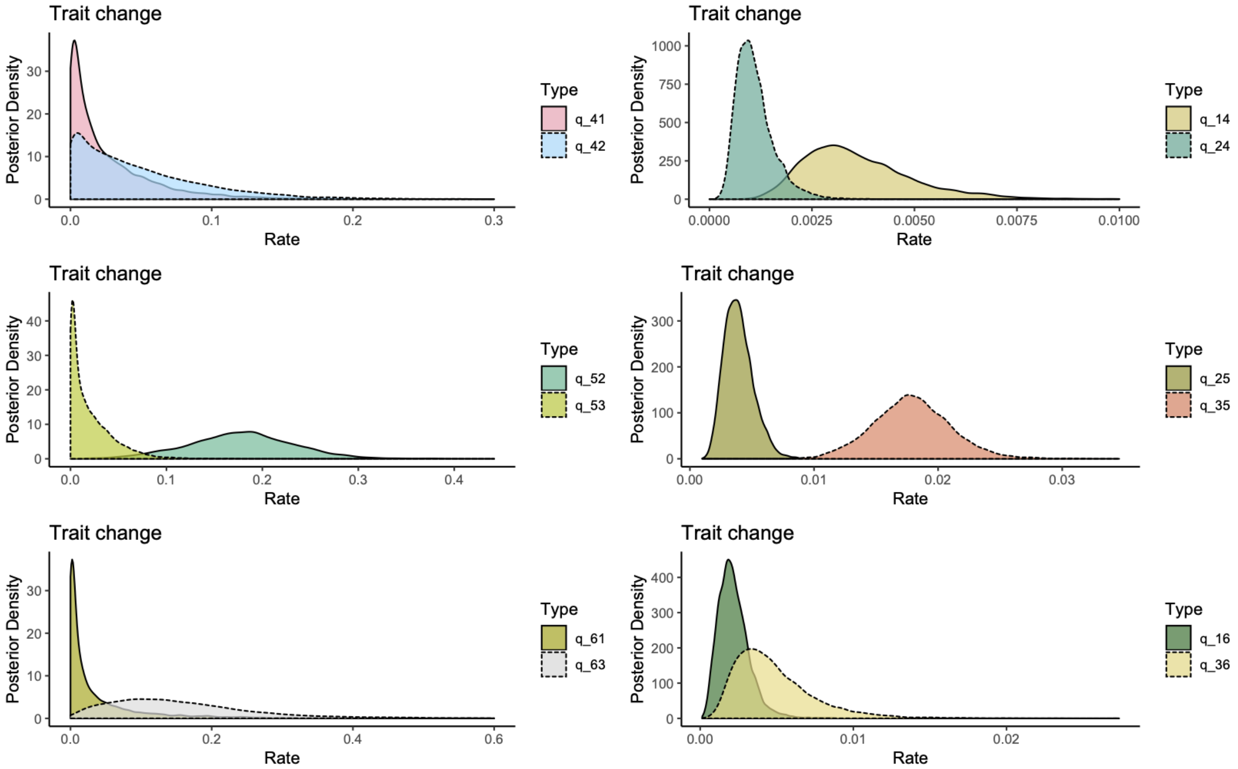


**Figure S8**. The posterior distributions of trait change for model MuSSE-6. The three panels on the first column show the transition rates out of the three states with two nest types (dome-cup, cup-hole, and dome-hole respectively) and the three panels on the second column indicate the transition rates towards the main states (dome-cup, cup-hole, and dome-hole respectively).

#### ***Full inference for MuHiSSE-6 model***

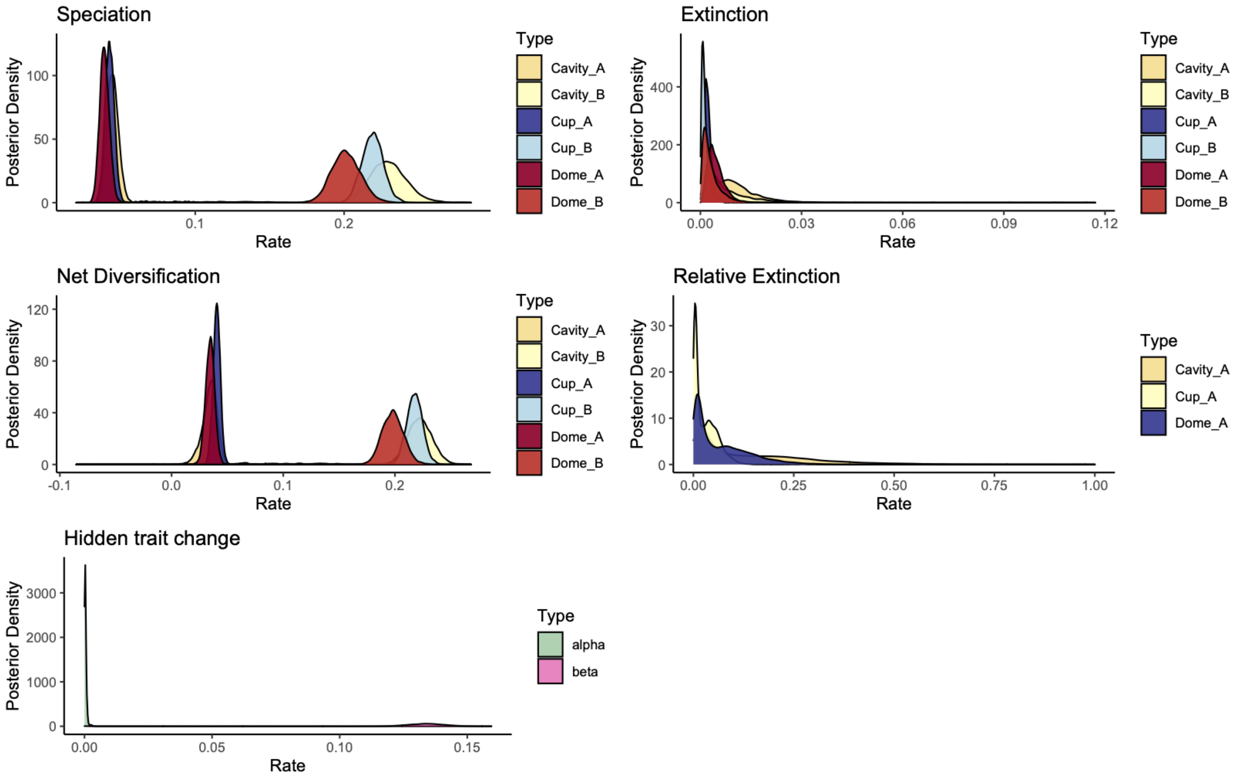


**Figure S9**. The posterior distributions for diversification parameters of MuSSE-6 model. The net diversification panel is represented as a violin plot in the main manuscript in figure 3b showing no differences in net diversification among the three states.

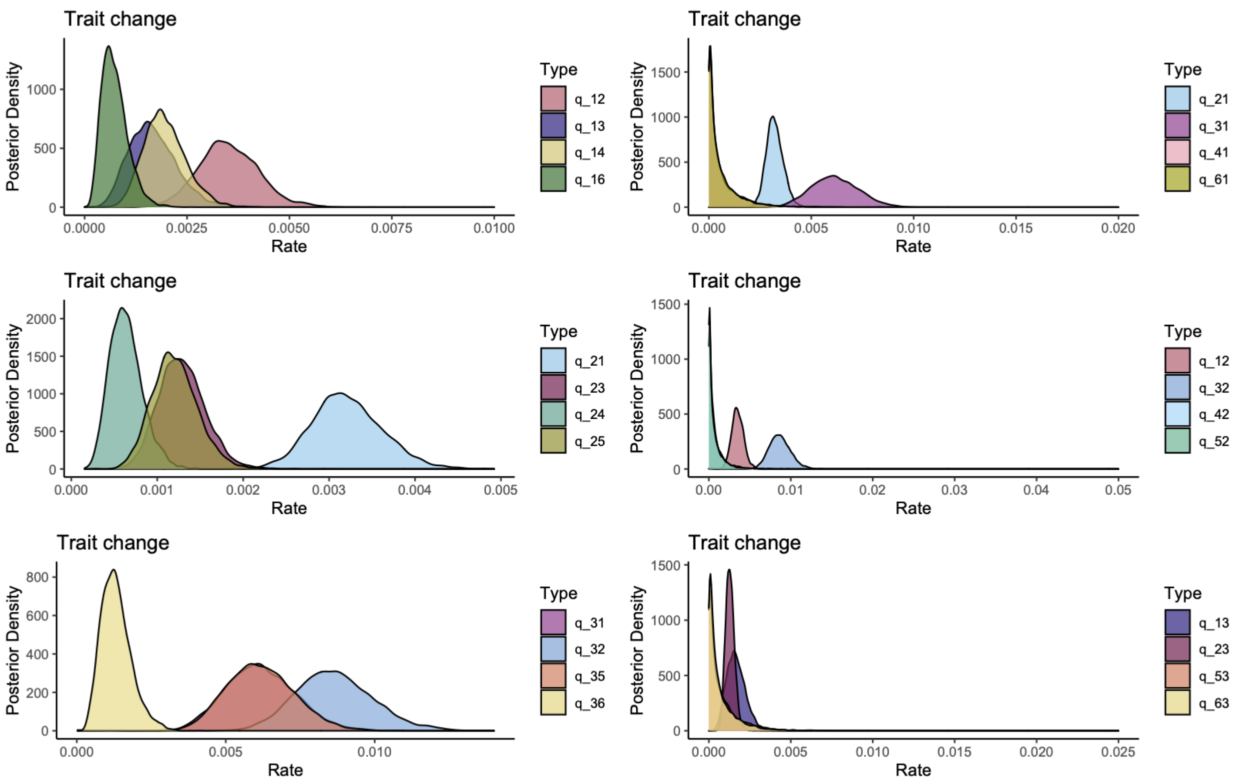


**Figure S10**. The posterior distributions of trait change for model MuHiSSE-6. The three panels on the first column show the transition rates out of the main three states (dome, cup, and hole respectively) and the three panels on the second column indicate the transition rates towards the main states (dome, cup, and cavity respectively).

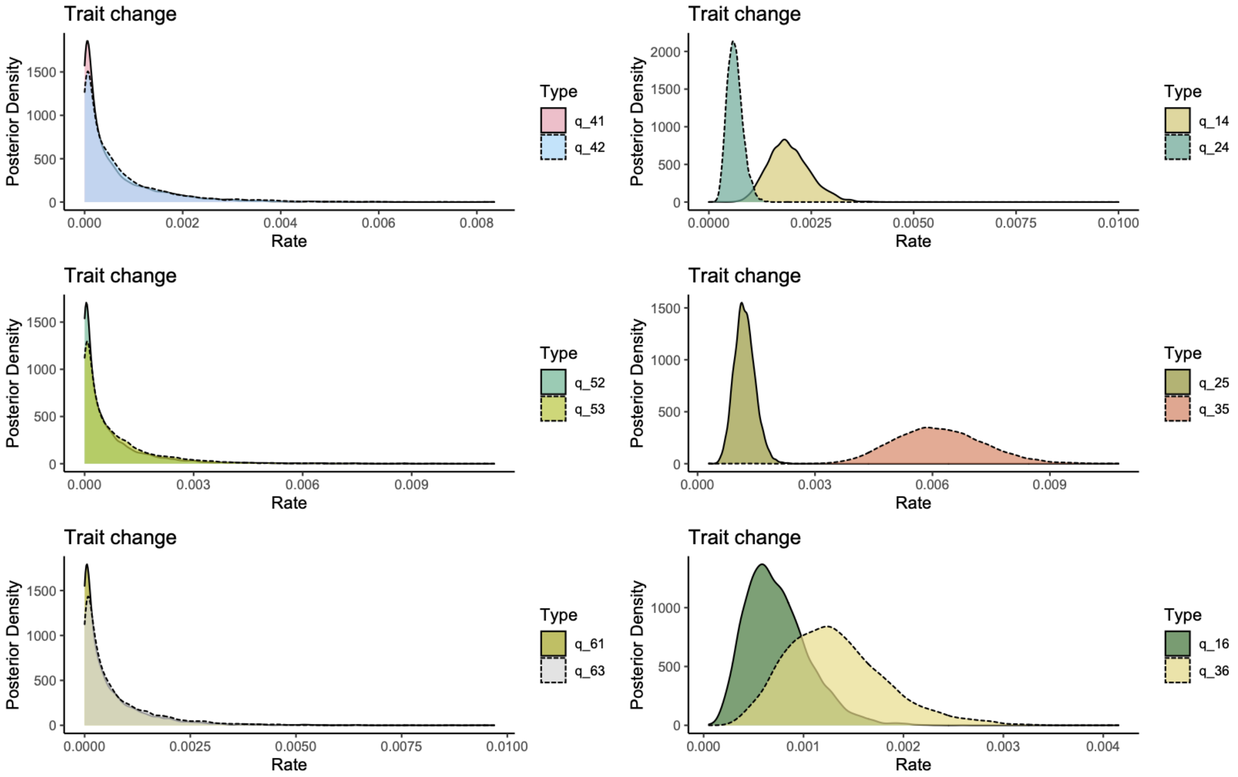


**Figure S11**. The posterior distributions of trait change for model MuHiSSE-6. The three panels on the first column show the transition rates out of the three states with two nest types (dome-cup, cup-hole, and dome-hole respectively) and the three panels on the second column indicate the transition rates towards the main states (dome-cup, cup-hole, and dome-hole respectively).

We did not calculate marginal log-likelihood for a CID-6 model to compare against the MuHiSSE-3 because CID-6 would require the estimation of too many parameters (36 in total: 12 diversification rates, 6 transition rates, 12 hidden state transition rates parameters, and 6 frequency parameters for the root), and, as discussed in the results, we found strong evidence that nest type is not linked to diversification using the much simpler CID-3 and the MuSSE-3. But visually comparing MuSSE-6 and MuHiSSE-6 we observe that there are no differences in diversification based on state, replicating the results of the 3 state models (Fig. 3).

**Full inference for MuSSE-3 model plus prior relative extinctions different for each state**

In order to measure the sensibility of the results due to the choice of priors of relative extinction in MuSSE-3 models, we defined a new model where the relative extinctions for dome, cup, and cavity were different and flipped. That is, we defined a prior for the relative extinction rate of state Hole with a minimum of 0.6, for state Cup with a minimum of 0.7, and state Dome with a minimum 0.8. The posterior distributions show similar results to the MuSSE with free estimation where Dome<Cup<Cavity in terms of net diversification but we observe more separation between the posterior distributions of net diversification per state. These results indicate that there is enough information in the data (through the likelihood function) so consistent results in the direction of the differences between states are found in the posterior distribution.

Assumptions of the MuSSE-3 with enhance extinctions are below:

| | | | |
|---|---|---|---|
| $\mu_C/\lambda_C \geq 0.8$<br>$\mu_D/\lambda_D \geq 0.7$<br>$\mu_H/\lambda_H \geq 0.6$ | $\mu_C = 0.8\lambda_C + \delta$<br>$\mu_D = 0.7\lambda_D + \delta$<br>$\mu_H = 0.6\lambda_H + \delta$ | $\lambda_i \sim LogN(\mu_\lambda, \sigma_\lambda)$<br>$\delta \sim Exp(2)$<br>$i = C, D, H$ | For cup, extinction rate is at least 0.8, for dome, is at least 0.7, and for hole, extinction rate is at least 0.6 of speciation rate. |

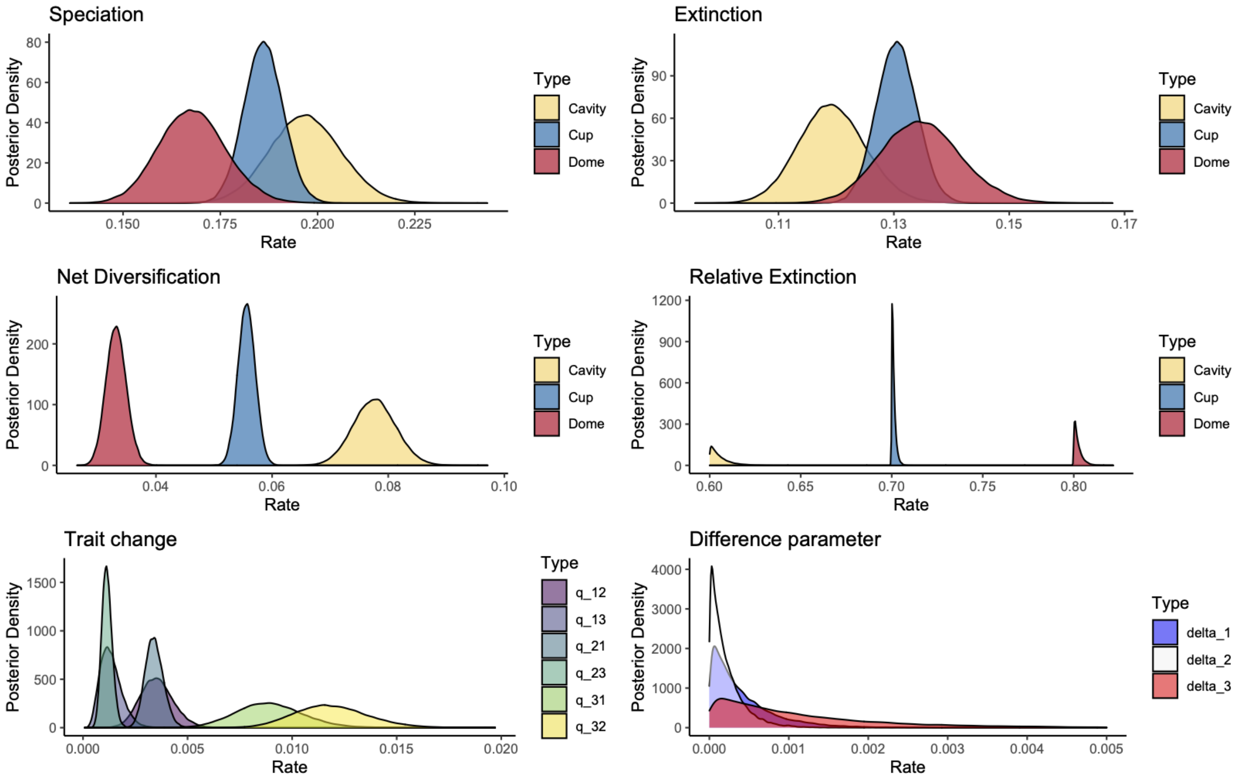


**Figure S12.** The posterior distributions for parameters in MuSSE-3+ prior relative extinction with a minimum of 0.6 for hole, 0.7 for cup, and 0.8 for dome. The different assumptions about relative extinction rates can be seen in the x-axis of the relative extinction panel. Net diversification differences between states conclude the same direction that the regular MuSSE (Dome<Cup<Hole).

#### Comment on eliciting prior distributions for faster rates of extinction

In our work, we chose to speed up extinction rates by utilizing the function $\mu = A\lambda + \delta$ which seems more complicated than using the more intuitive function $\mu = \lambda * \delta$. In our first attempt, we fitted the latter and more intuitive approach. However, diversification models have a strong tendency to underestimate extinction rates due to the lack of extinction observations in incomplete phylogenies (Louca and Pennell (2021) has the exact mathematical demonstration of this phenomenon). Therefore, when fitting $\mu = \lambda * \delta$, the model estimation return ssimply the unconstrained results (same as the results of the simple MuSSE-3) but rescaled by the factor 1/ $\delta$ failing at setting a lower bound of extinction. By forcing the lower bound to be a fixed proportion A in the function $\mu = A\lambda + \delta$, and allowing for some stochasticity with random variable $\delta$ it is possible to restrict parametric space successfully and force estimations of extinction that are larger or equal than the set lower bound, which is the scenario we were interested in.

**Ancestral state reconstruction**

When using phylogenetic tree with passerines only, the maximum a posteriori for the root of the clade has probability 0.51 for dome, 0.41 for hole nesting, and 0.08 for cup nesting (Fig.S13).

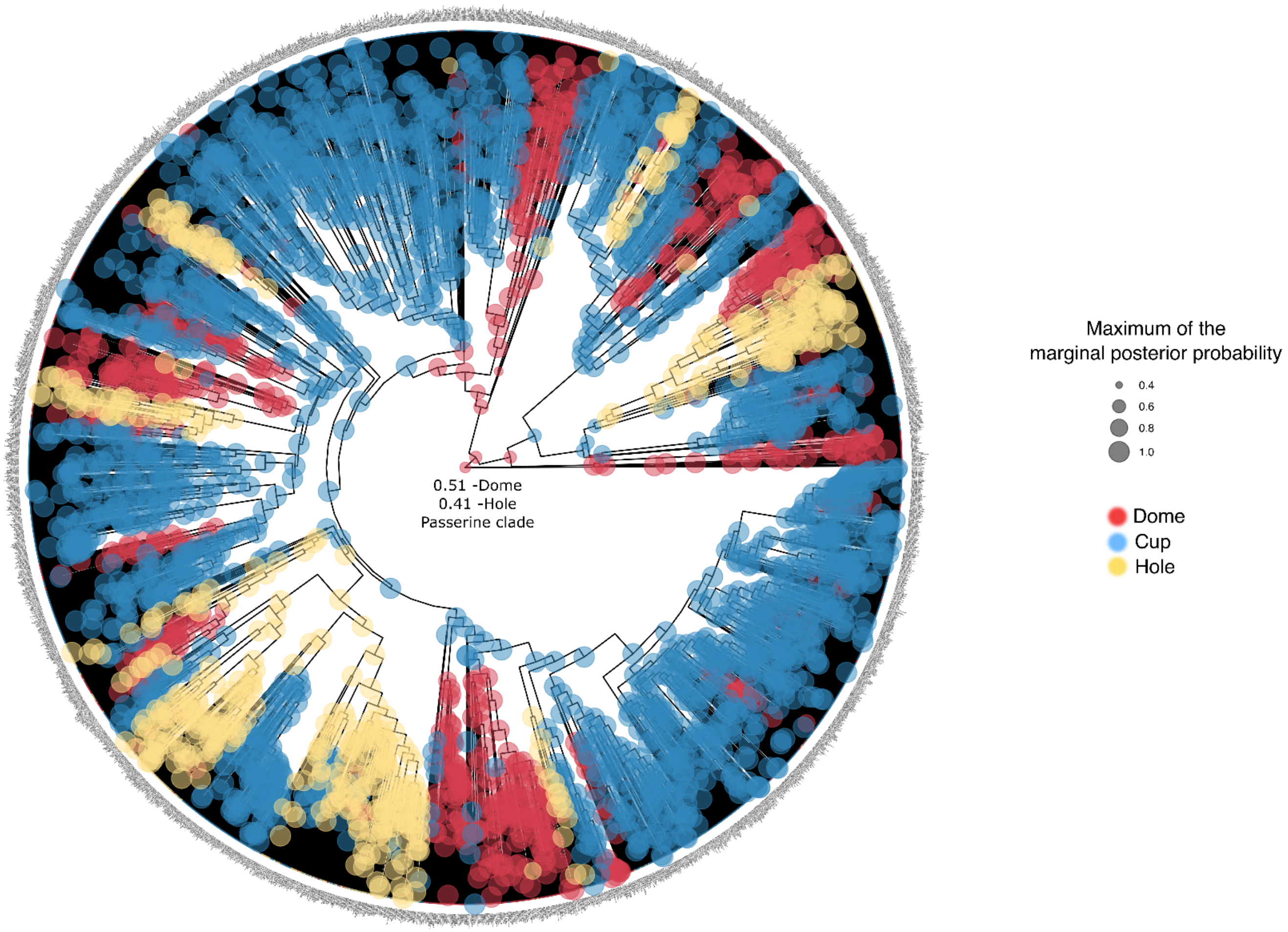


**Figure S13.** Ancestral state reconstruction of nest type using passerine lineages only undear a Markov model with three states (Mk3). The size of a node represents the maximum probability value of the marginal posterior distribution of the node. The color of the node represents the nest type associated with the maximum posterior probability. Figure made using RevGadgets (Tribble et al. 2022)

**Eliciting prior distributions and impact of hyperparameters**

For all the diversification analyses in the main manuscript we chose to model the log-transformed speciation and extinction rates using a normal distribution for each of the parameters. Each of the normal distributions have an expected mean value $\eta_\lambda$=-2.16 which represents the natural log of natural the log of 3225 divided by two, following with a second division by the clade age that is 64.5 my. The standard deviation parameter $\nu_{\boldsymbol{\lambda}}$is equal to 1.12.

That is, the probability density function for the log-transformed speciation rate $ln(\lambda)$ is

$$ln(\lambda) \sim N\left(\eta_\lambda = ln\left(ln\frac{(3225/2)}{64.5}\right) =- 2.16\,, \nu_{\boldsymbol{\lambda}} = 1.12\right).$$

These choices of the prior distributions and hyperparameter imply that the mean for speciation rates $\lambda$ is a log-Normal distribution with approximately $\boldsymbol{\mu}_{\boldsymbol{\lambda}} = e^{\eta_\lambda} = e^{-2.167} = 0.11$ (interpreted as 0.11 lineages splitting per my). Notice we calculated the approximate transformation of the mean since the true transformation from the log-normal to the normal would require that the mean is calculated using the standard deviation $\nu_\lambda$. The choice of standard deviation for the log-transformed speciation rate also implies that the standard deviation for the untransformed speciation rate $\lambda$ is $\sigma_\lambda = 3.3$.

In mathematical terms the prior distribution of the speciation (and extinction) rates in our models are approximately

$$\lambda \sim logN(\mu_\lambda = 0.11,\ \sigma_\lambda = 3.3).$$

The log-normal distributions assure that all speciation and extinction rates will be positive but allow for broad uncertainty on diversification rates a priori. In figure S14 we are illustrating this prior distribution for one speciation rate.

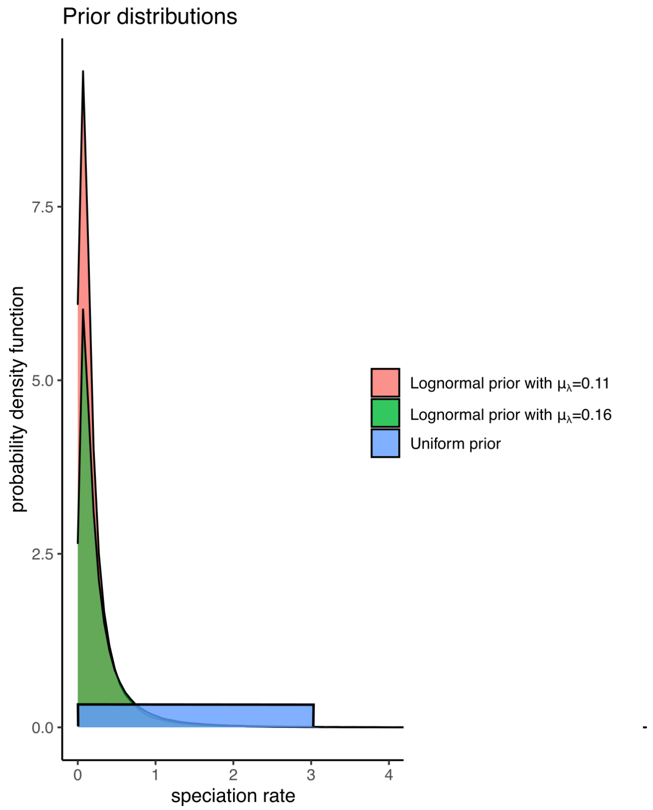


**Figure S14.** We tested three different prior distributions for MuSSE-3 to test for the sensitivity of the results. The first two distributions are lognormal with different means, one calculated from extant passerine lineages with nest type data with an older tree, and the second calculated using all extant passerine taxa with a crown age of 50 my. The third distribution naively assumes that speciation and extinction rates are between 0 and 3 lineage changes per million year.

As priors are elicited subjectively, we can consider alternative scenarios for deciding values. For example, if we consider the whole diversity of the passerine clade with about 6,500 species, the speciation rate average of choice in Passeriformes would be $\mu_\lambda =$ log(6,500/2)/64.5 = 0.13. We can also consider a different crown age like 50 my without the *Strigops* outgroup that is considered in our main manuscript), in which case the rate would be $\mu_\lambda$=0.16. This exercise is extremely valuable to evaluate the sensitivity of the inference due to prior distribution selection but most importantly we would know if the likelihood function in a Bayesian analysis is informative. This is the case in our work, as we show in Fig. S15 different choices of priors for speciation result in the same net diversification rate inference for our three states.

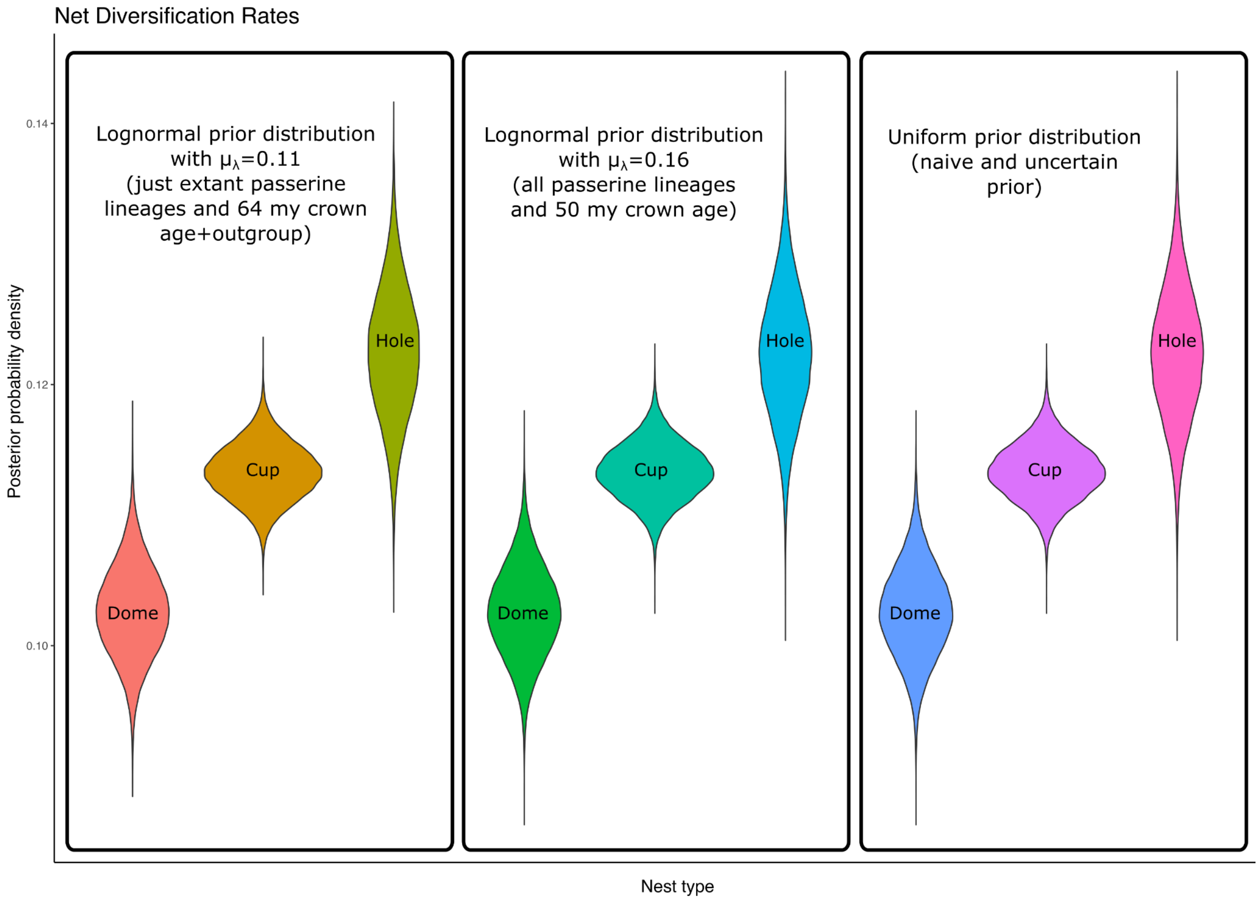


**Figure S15.** Estimates of net diversification per nest type under three different prior assumptions. From left to right: The first panel assumed all rates of speciation and extinction a priori are log-normal with a mean of 0.11 as presented in the main manuscript. The middle panel assumed all rates of speciation and extinction a priori are log-normal with a mean of 0.16. The third panel assumed all rates of speciation and extinction are Uniform(0,3). The inference is consistent across prior assumptions for all net diversification, showing that the likelihood from our example is informative.
